# Climate-change risk analysis for global urban forests

**DOI:** 10.1101/2021.05.09.443030

**Authors:** Manuel Esperon-Rodriguez, John B. Baumgartner, Linda J. Beaumont, Jonathan Lenoir, David Nipperess, Sally A. Power, Benoît Richard, Paul D. Rymer, Mark G. Tjoelker, Rachael V. Gallagher

## Abstract

Urban forests (i.e. all vegetation present in urban areas), provide environmental and socioeconomic benefits^1^ to more than half of the global population^2^. Projected climate change threatens these benefits to society^3–5^. Here, we assess vulnerability to climate change of 16,006 plant species present in the urban forests of 1,010 cities within 93 countries, using three vulnerability metrics: exposure, safety margin and risk. Exposure expresses the magnitude of projected changes in climate in a given area, safety margin measures species’ sensitivity to climate change, and risk is the difference between exposure and safety margin^6^. We identified 9,676 (60.5%) and 8,344 (52.1%) species exceeding their current climatic tolerance (i.e. safety margin) for mean annual temperature (MAT) and annual precipitation (AP), respectively. By 2050, 13,479 (84.2%) and 9,960 (62.2%) species are predicted to be at risk from projected changes in MAT and AP, respectively, with risk increasing in cities at lower latitudes. Our results can aid evaluation of the impacts of climate change on urban forests and identify the species most at risk. Considering future climates when selecting species for urban plantings will enhance the long-term societal benefits provided by urban forests, including their contribution to mitigating the magnitude and impacts of climate change.

## Introduction

Urban areas span approximately three percent of the Earth’s land surface area^7^, and accommodate more than 4.2 billion people, representing 55% of the global population^2^. Within cities, vegetation present in parks, woodlands, abandoned sites and residential areas and along streets - collectively referred to as urban forests^8^ - provide environmental services and socioeconomic benefits, such as carbon sequestration and heat mitigation throughout microclimate processes^1^. Furthermore, urban forests are crucial for mental health improvement in time of societal crisis, such as the COVID-19 pandemic^9^. By 2050, cities are expected to expand in size and in population around the globe, with predictions of 6.6 billion people living in urban areas by this time (~70% of the predicted global population)^2^. As human population grows, so too will the societal demand on urban forests.

The pace at which climate is changing^10^ poses a serious threat to the continued persistence of urban forests globally. While the impacts of climate change on natural ecosystems, both terrestrial and aquatic, have been widely studied^11–13^, less is known about potential impacts on urban ecosystems (but see^14,15^). To date, there have been no global assessments of the vulnerability of urban forests to future climate change, although several independent regional-scale studies have been conducted^3,5^. Urban forests have the potential to mitigate the adverse effects of global climate change by shading buildings and paved surfaces to reduce energy usage^16^, decreasing urban heat through evapotranspiration, capturing greenhouse gases and storing carbon trough photosynthesis^17^, thereby collectively contributing to limiting the rise in global temperature to 1.5°C above pre-industrial levels as outlined in the Paris Agreement^18^. Therefore, there are good reasons to assume that planting and preserving climate-resilient plant species, in public and private space in cities, can contribute to mitigating the adverse impacts of global climate change, and enable urban forests to continue to play an essential role in people connection to nature^1^.

Natural and urban ecosystems are already being impacted by changing climate conditions resulting in sub-optimal growth and increased mortality^19,20^. Climate change exacerbates the risk of severe summer drought in many regions of the world and extreme events – such as heatwaves and cyclones, which are predicted to increase in frequency and severity^21,22^ – are already contributing to extensive tree mortality globally^19,23^ (**Figure 1**). In addition, characteristics of the urban environment, including high cover of impervious surfaces and the urban heat island effect, can locally exacerbate climatic extremes^24^. Species mortality in cities, therefore, can have environmental and socio-economic consequences for local governments and urban residents^23,25^. For instance, a study of urban areas in the USA estimated that a decrease of 1% in urban tree cover may represent economic losses of USD ~$100 million per year due to the associated loss of air pollution control and carbon sequestration, and increased energy consumption and power plant emissions^20^.

**Figure 1.**
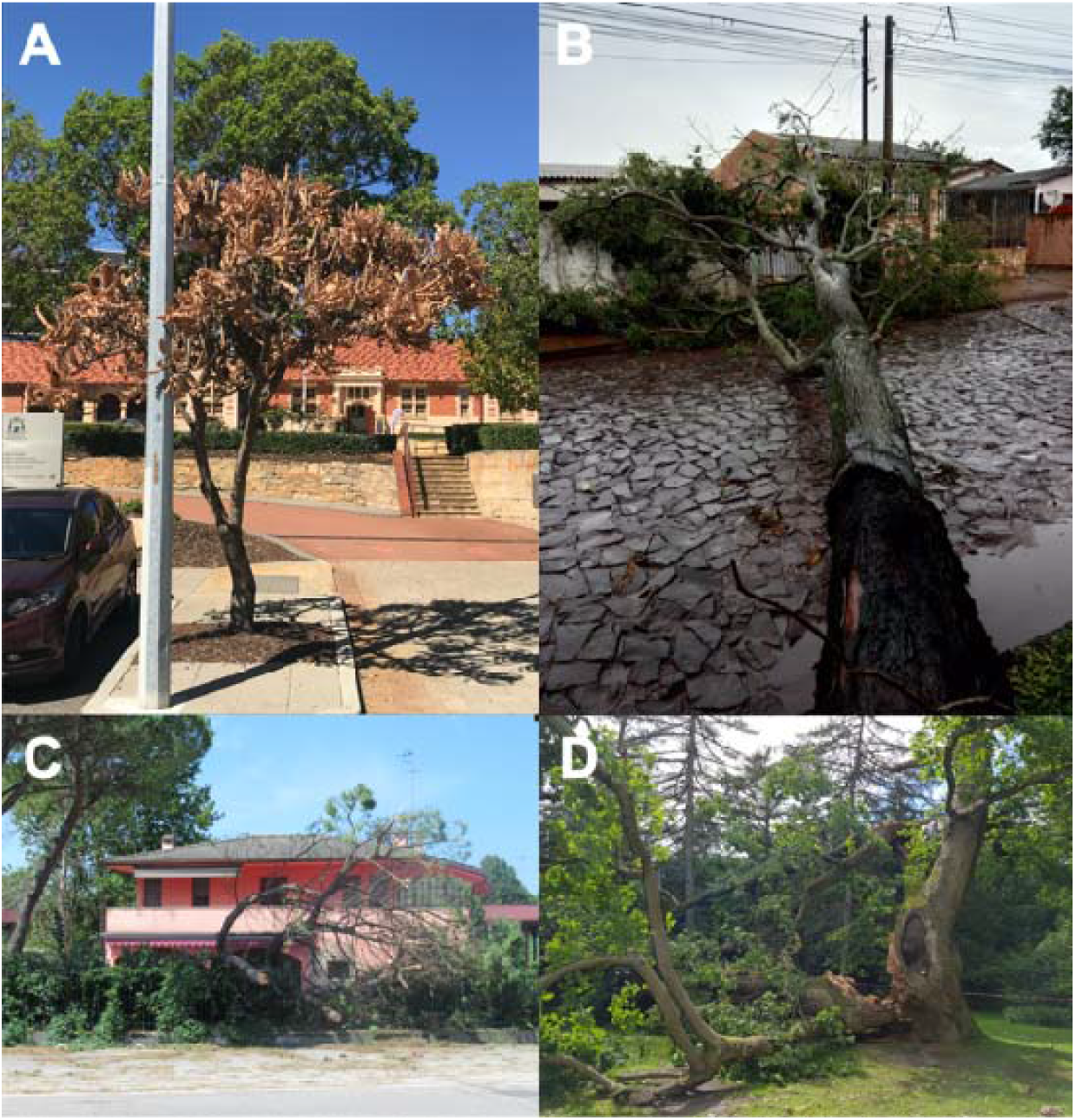
Examples of urban tree mortality as a result of extreme weather events globally: (a) Banksia spp. dieback after an extreme heat and drought event in Perth, Australia; (b) tree uprooted by wind and storm in Foz do Iguaçu, Brazil; (c) tree damage associated with a cyclone in Padua, Italy; and (d) storm damage to an oak tree in Alnarp, Sweden. Photos provided by MER, Evaldo Monteiro Guimarães, Alessio Russo and Johan Östberg.

Strategically, given the slow growth rates of trees and the importance of promoting tree longevity, successful urban greening involving targeted increases in tree canopy cover must be planned with future climatic conditions in mind for the next decades to secure the persistence of urban forests into the future. Urban greening is in part directed towards the strategic delivery of ecosystem services and benefits^26^. Global estimates of the value of urban forests exceed USD $500,000,000 per annum, as the sum of the services urban forests provide to society^27^. To deliver these services effectively in the long term, it is crucial to identify vulnerable species and quantify their risk of mortality under future climates. Policymakers and urban forest managers require to optimise the use of resources and minimise losses when investing in urban forestry programs^25^. Identifying species and cities most at risk from climate change is crucial for ensuring urban forests are made resilient in the long term.

Here, we present a climate-risk analysis for global urban forests. We assessed the potential impacts of future changes in climate for 16,006 vascular plant species from 1,010 cities in 93 countries (**Figure 2**). To do so, we calculated three metrics: (1) exposure, the magnitude of projected changes in temperature and precipitation (assessed using the 5^th^ and 95^th^ percentiles of temperature and precipitation variables, respectively; see details in Methods) at each city; (2) safety margin, the sensitivity of each species to changes in climate according to its climatic tolerance; and (3) risk, calculated as the difference between exposure and safety margin^4,6^. We calculated a species’ climatic tolerance from its current geographical distribution, while its climate risk was determined by the projected future climate in cities where the species is currently planted and growing. Because of the mismatch and asynchrony between the speed at which contemporary climate change is happening and the time required for long-lived plant species (e.g. trees and shrubs) to respond to climate change^28,29^, aka the climatic debt^30^, we assume that a significant proportion of species in urban areas are already at risk or partially decoupled from macroclimatic conditions thanks to costly management practices (e.g. water supply). Hence, contemporary urban planning and species selection require to ensure a successful climate mitigation strategy for the future.

**Figure 2.**
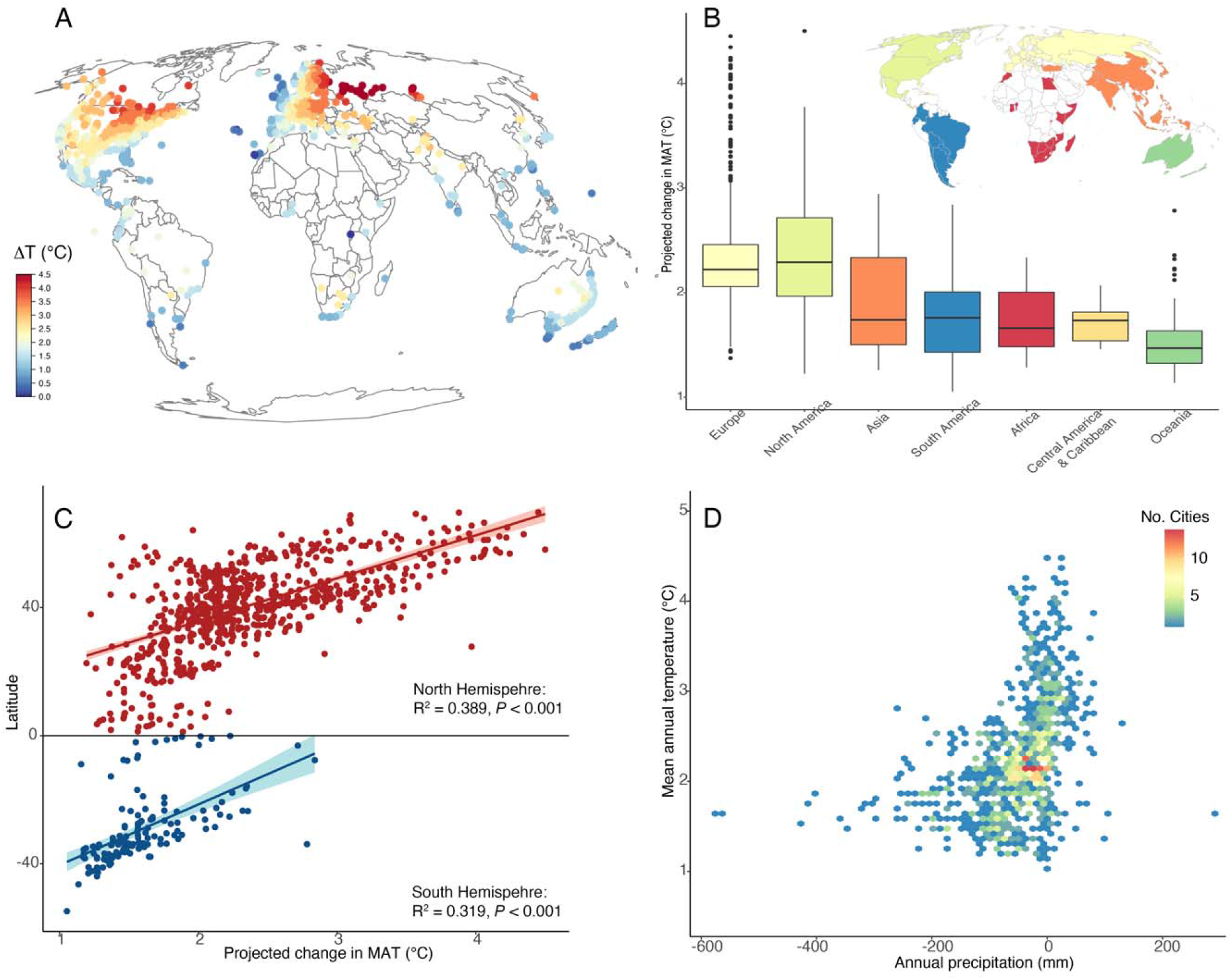
Exposure to climate change across the world’s cities. Exposure of 1,010 cities to changes in mean annual temperature by 2050 (MAT; A) and boxplot of changes in MAT in seven geographical regions (Appendix 1; B); exposure to temperature change across the latitudinal gradient of cities in the Northern and Southern Hemisphere with 95% confidence interval for predictions from a linear model [“lm”]) (C); and the interaction between exposure to changes in MAT and annual precipitation (mm) in 2050 (D). All plots display data for RCP 6.0.

## Results

### Exposure

Exposure (*E*) measures the magnitude of change in climate conditions in a given city between current (baseline average during 1979-2013) and future (2050 or 2070) climatic conditions. Under the Representative Concentration Pathway (RCP) 6.0 (for RCP 4.5, see **Table S1; Figure S1**) and according to an ensemble of 10 General Circulation Models (GCMs; see details in Methods), mean annual temperature (MAT) across all 1,010 cities in this study is predicted to increase by an average of 2.3°C (standard deviation ± 0.6°C) by 2050. The greatest increases are predicted to occur in cities at higher latitudes in the Northern Hemisphere. The increase in MAT is predicted to exceed 3°C for 126 cities (**Figure 2**). By 2070, average MAT is predicted to be 3.2°C (± 0.9°C) warmer than baseline conditions under the same RCP and set of GCMs, with 570 cities predicted to experience >3°C of warming. For annual precipitation (AP), 820 cities are predicted to become drier in 2050 with declines averaging −68 mm (± 68.4 mm). By 2070, AP declines −81.7 mm (± 74.7 mm) in 805 cities (**Table S1; Figure S2**). See Supplemental material for details on changes (i.e. exposure) of the maximum temperature of the warmest month (MTWM) and precipitation of the driest quarter (PDQ).

### Safety margin

The safety margin (*S*) describes a species’ potential tolerance to changing climate conditions within a given city. For each climate variable (i.e. MAT, AP, MTWM, PDQ), *S* is calculated under current climate as the difference between the value of that variable for the city and the species’ tolerance limit for that variable, estimated as the 5^th^ percentile for precipitation and 95^th^ percentile for temperature across the entire geographic range of the species (**Figure S3**). We note that the realised niches for many species are likely underestimating their resilience to climate change given our data comes from occurrence records (see details in Methods), where biotic interactions may restrict the realised niche limits and/or sampling bias may affect estimates. Species may indeed tolerate hotter or drier condition than what they experience in the field.

The safety margin indicates how much warmer (or drier), a city could become in the future before a species currently planted within the city exceeds the limits of its realised climatic niche^4,6^. However, it is possible that some species are already at risk under current climatic conditions with today’s temperature being already warmer or precipitation lower than the threshold the focal species can withstand (**Figure S3**). We found that 9,676 species (60.5%) are presently exceeding their MAT safety margin (i.e. “unsafe”) in at least one city where they are planted (**Figure S4**). Three cities were identified as currently having 100% of their species exceeding their MAT safety margin (Chennai and Pondicherry, India; Manila, Philippines). Furthermore, 86 cities were identified as having >50% of their species exceeding their MAT safety margin (**Figure S5**).

For AP, 8,344 species (52.1%) are already exceeding their AP safety margin in at least one city where they are planted. Three cities were identified as having 100% of their species exceeding their AP safety margin (Ankara, Turkey; Gilgit and Quetta, Pakistan), while 89 cities had >50% of their species currently exceeding their AP safety margin (**Figure S5**). See Supplemental material for details on species’ safety margin of MTWM and PDQ.

### Risk

Risk (R) to urban forests under future climates is defined as the difference between the city’s exposure to future climate change and its species’ current safety margin (*R* = *E* – S)^6^. By 2050, we found that increasing MAT under RCP 6.0 may result in 13,479 species (84.2%) being at risk (i.e. city’s future climate will exceed the realised climatic tolerance of species; S: *R* > 0) in at least one city where they are currently planted (for RCP 4.5, see **Figure S6**). Although most cities threatened by future increases in MAT are concentrated in the USA (320 cities), Australia (86 cities), Mexico (46 cities), Germany (45 cities), France (43 cities), the UK (43 cities) and Canada (35 cities), we found a tendency for the risk to increase towards the Equator (**Figures 3**–**4**). For a subset of 37 cities, the exposure level is predicted to be so high under future conditions (by 2050, RCP 6.0) that it will represent a risk to all (100%) of the recorded species that currently occur in these cities. By 2070, 14,209 species (88.8%) are predicted to become at risk (**Figure S6**), with 51 and 602 cities predicted to have 100% and >50% of their species at risk, respectively.

**Figure 3.**
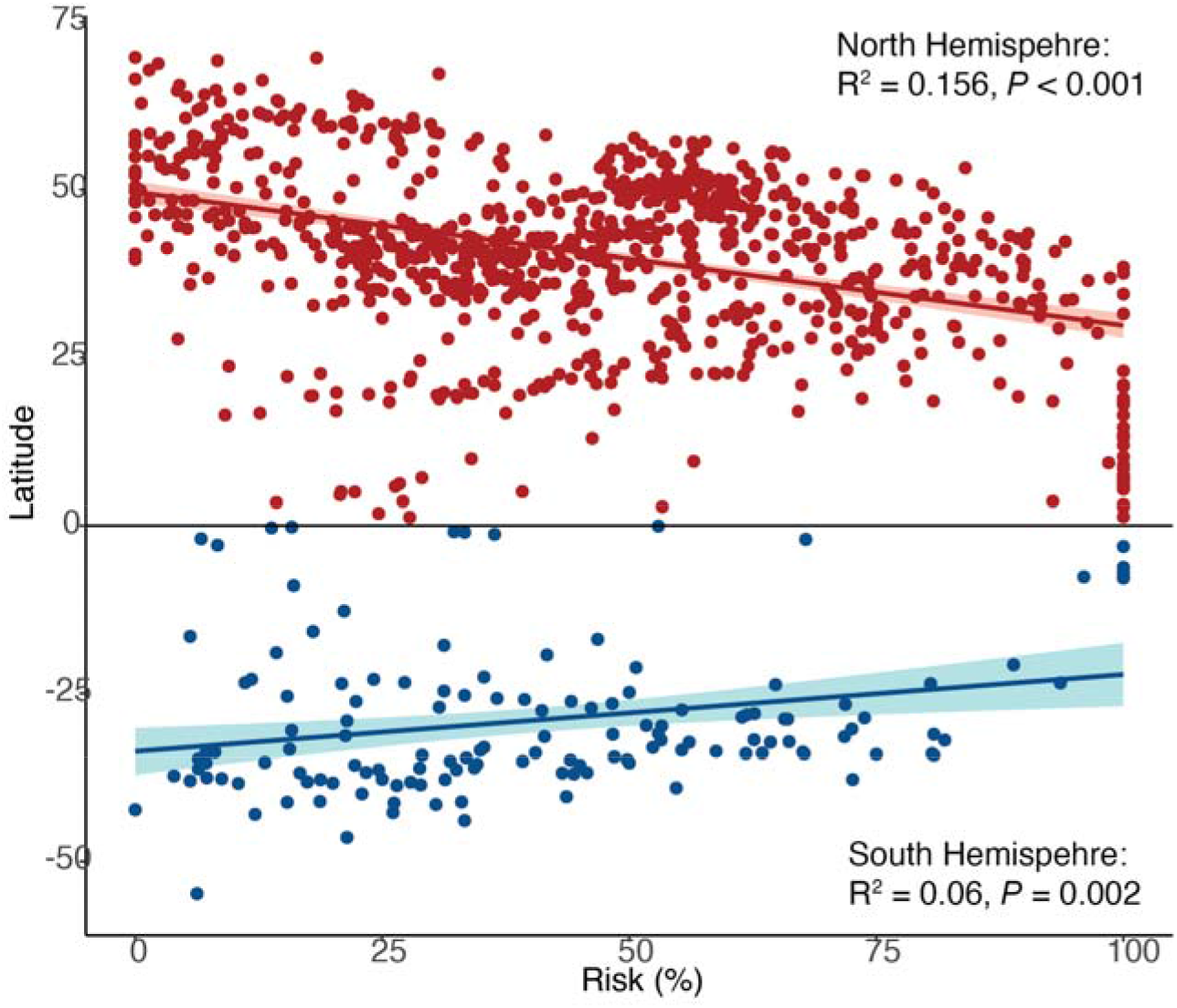
Relationship between risk and latitude (n = 870 cities in the northern hemisphere and 140 in the southern hemisphere). Ribbons indicate the 95% confidence interval for predictions from a linear model [“lm”]. Each point represents the mean risk for all species in a given city based on MAT exposure and safety margin. Data for 2050 and RCP6.0.

**Figure 4.**
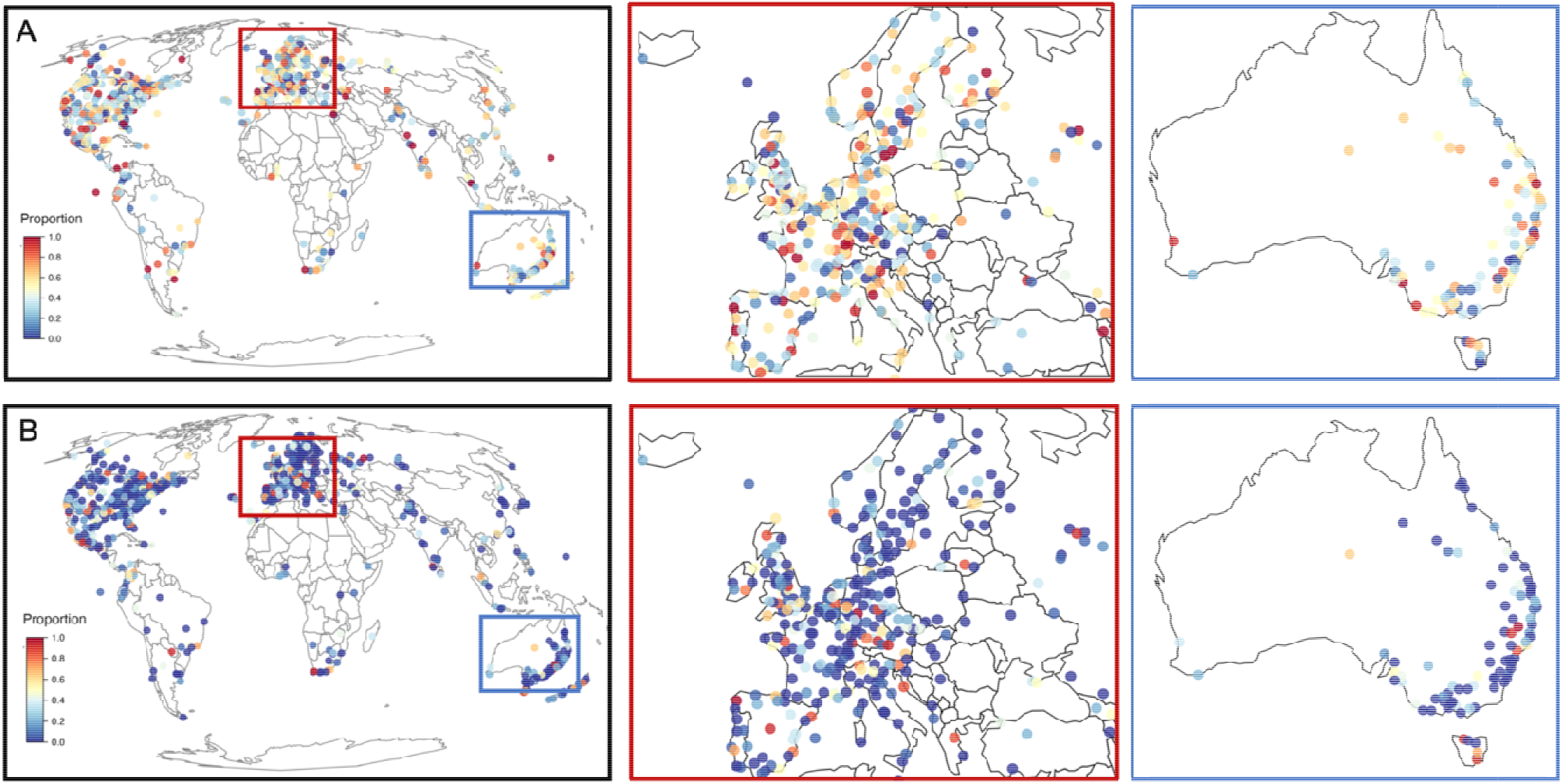
Proportion of urban forest species in each city predicted to be at risk from projected changes in mean annual temperature (A; MAT) and annual precipitation (B; AP). Data for 2050 and RCP6.0.

Under future AP conditions, 9,960 species (62.2%) are predicted to be at risk (i.e. drier conditions than the focal species’ realised niche limit) in at least one city where they are currently planted. We identified five and 136 cities with 100% and >50% of their species at risk, with the highest concentrations of cities found in the USA (257 cities), Australia (77 cities), Mexico (42 cities), Germany (37 cities), France (32 cities), Russia (30 cities) and the UK (30 cities) (**Figure 4**). By 2070, 10,233 species (64%) are predicted to become at risk of decreases in AP, with five and 151 cities having 100% and >50% of their species at risk, respectively. See Supplemental material for details on species’ risk of MTWM and PDQ and **Figures S7**–**S11**.

## Discussion

We show that 1,010 and 820 cities are at risk from MAT and AP change by 2050. Climate-driven changes to urban forests will have adverse consequences for city dwellers and governments globally, although the magnitude of these consequences will vary across cities. Cities predicted to have higher increases in MAT (>4 °C) are more vulnerable to climate change. Further, cities’ vulnerability can be exacerbated by the urban heat island effect and population size^31^.

Changes in precipitation patterns, which are likely to shape species’ growth and survival, will also impact urban forests globally. In general, urban forests that experience declines in precipitation will be more vulnerable than those facing higher rainfall, although significant increases in AP might also represent a risk factor, i.e. flooding^32^. Human management such as irrigation or stormwater capture can aid at mitigating the adverse effects of low precipitation by providing Supplemental water during periods of severe climate stress^33^ and promoting evapotranspiration (local cooling effect generated by plants), which will be crucial to mitigate future heatwaves in cities^34^. Irrigation, nonetheless, may not be a sustainable solution in many places where water is increasingly scarce^35,36^. This type of costly management practices may actually explain why so many species are already exceeding their current AP safety margins.

Urban forests are often water stressed or closely coupled to regional precipitation and water balance; hence, species growing under hydrologically stressful conditions are more vulnerable to extreme climate events^14^, resulting in higher mortality rates^37^. Unfortunately, studies of urban species mortality driven by climate change are rare and, generally, anecdotal and limited in scope and broad applicability^38^, which limits the capacity to assess climate risk for those species that are currently experiencing conditions that exceed their safety margins^4^. However, our results show that some cities currently harbor many species living outside of their realised climatic tolerance. We found a high number of species currently exceeding their safety margin for the four climate variables (MAT = 60.5%; AP = 52.1%; MTWM = 56.3%; PDQ = 46.8%), suggesting there are additional management actions (e.g. irrigation) and biological factors facilitating species’ presence in cities and decoupling them from macroclimatic fluctuations. Being planted in an area, however, does not necessarily mean that a species is performing well in that location. There is also a difference between being able to tolerate and persist in certain conditions and being able to maintain a function. That is, species whose safety margins are exceeded may be able to tolerate, but would not have the capacity to function and remain healthy under those conditions. The long-term stability of urban forests, therefore, depends on the identification of species and cultivars that are resilient to long-term climate change in a given location and are able to thrive and survive^39^.

By 2050, the number of species at risk is predicted to increase for all climate variables except for AP, where predicted rainfall increases in some cities might ameliorate the impacts of climate change. However, seasonal decreases in rainfall (i.e. PDQ) increase the risk that a given species experiences conditions that are drier than it can tolerate – as indicated by its safety margin. Further, global warming increases risk in cities toward lower latitudes (**Figure 3**), where resources to mitigate climate change are more limited^40^. In these cities, MAT exposure is lower (particularly in the Northern hemisphere) compare to cities at higher latitudes; therefore, species’ safety margin might be driving the increase in risk. This highlights a potential mismatch between species selection in those cities and the changing climatic conditions that have occurred during over last decades (i.e. baseline 1970-2013).

We also found 12,653 (79.1%) species at risk from increases in MTWM, highlighting that extreme temperatures represent a significant threat to urban forests, especially towards lower latitudes. Predicted changes in extreme/seasonal variables (i.e. MTWM, PDQ), therefore, impose a thermal/hydrological stress to plant species, as well as human populations. Our risk approach allows identification of the most vulnerable urban plant species^4,6^. Although here we described risk in a binary manner (i.e. high/low risk), our risk estimation can provide details on which species are more at risk of changes in climate (**Figure S12**) and can guide prioritization and substitution for more resilient species.

Climate change will become a key driver of species mortality in urban forests^14^. Likely, it will become increasingly more difficult to mitigate the effects of climate change through management actions, such as irrigation, to offset soil water deficits, particularly under limited urban water supply. Additionally, management options for altering or mitigating rising temperatures, particularly maxima and minima, are limited^41^. Furthermore, vulnerable species will require more intensive and costly management actions and in extreme cases, replacement, if they cannot cope with climate change. Our results, therefore, provide a path forward to better inform local governments of potential risks and improve species selection given future climates, to avoid planting failures and maximise societal benefits in a future warmer world. However, the mitigation of climate change impacts ultimately will depend on the available resources of each city and its capacity to respond and cope with climatic changes as they occur.

To improve species selection in relation to climate, however, requires studies of realised niche matching (i.e. species’ niche with city’s climate), physiological plasticity and environmental tolerance of urban species and cultivars. Selection of resilient species should be based on life history and physiological information. Currently, these data are limited, which increases uncertainty around decision-making^42^. To maintain healthy urban forests in a changing climate, it will be necessary to address budget considerations, the provision of adequate time and effort for establishing and maintaining urban plantings, and filling the knowledge gaps in appropriate species selection for changing climatic conditions. We emphasize the importance of taking immediate actions to secure the survival and persistence of urban forests globally and avoid the collapse of these socio-ecosystems.

## Methods

### Urban forest composition and urban areas

Although urban forests are mainly defined as systems comprising trees^43^, this definition excludes the ecosystem services provided by other plants’ growth forms. Therefore, we adopted a broader definition, which considers an urban forest as all vegetation (i.e. tracheophytes: vascular plants) present in urban parks, woodland, abandoned sites, residential areas, private gardens and along urban streets (sensu ^8^).

We obtained occurrence records globally for all tracheophytes from the Global Biodiversity Information Facility (i.e. 181,914,869 records from 3,949 published datasets) (GBIF.org; 18 December 2019 GBIF Occurrence Download https://doi.org/10.15468/dl.cpwlwc). We only retained occurrence records with sufficient information on geographical coordinates. Additionally, occurrence records were filtered and cleaned by removing spatially invalid or suspect records that could lead to miscalculation of species’ climate niches and duplicate records using the CoordinateCleaner package ^44^ in *R* version 4.0.5^45^. Finally, in order to refine the list of species to be included in the subsequent analyses, we removed all species with no occurrence records located inside the geographical boundaries of urban areas (see below). This resulted in 23,857,682 valid occurrence records, from 16,006 species within 342 families. The average number of species per family was 48 (± 148 species), with a maximum of 1,860 species in Asteraceae, while 64 families were represented by a single species. Taxonomy was standardized and verified against GBIF and then against The Plant List (TPL; www.theplantlist.org) using Taxonstand, taxize, and taxizehelper packages^46–48^ in R^45^.

Polygons defining the boundaries of 6,018 urban areas (i.e. cities) globally were obtained from Kelso and Patterson ^49^ as a shapefile (WGS84; 1:10 million; EPSG:4326). These data were projected to the Mollweide projection, an equal-area pseudocylindrical map projection (ESRI:54009).

We assessed the effect of potentially inadequate sampling on our analyses by estimating the completeness of the species inventory in each city (see details in **Supplemental Material**). Based on this analysis, we retained 1,010 cities from 93 countries with high completeness (>0.20). The average number of cities within countries was 11 (± 35 cities), with a maximum of 324 in the USA, while 44 countries were represented by a single city. The average number of species per city was 217 (± 294 species), with a maximum of 2,249 in Brussels (Belgium) and a minimum of 11 species in ten cities from Albania and Palestine (**Figures S13**–**S14**).

### Climate data

Baseline and future climate data were obtained from CHELSA Version 1.2 (climatologies at high resolution for the earth’s land surface areas^50^) at a spatial resolution of 30 arc-seconds (∼1 km at the equator). A detailed description of the generation of these data is given elsewhere^50^. We selected four climate variables; two of them describing mean conditions: (1) mean annual temperature (MAT) and (2) annual precipitation (AP), and two variables describing extremes of climate: (3) maximum temperature of the warmest month (MTWM) and (4) precipitation of the driest quarter (PDQ) (**Table S2**). All climate data were projected to the Mollweide projection system (ESRI:54009) at a 1 km resolution using bilinear interpolation. Baseline data represent climate conditions during the period 1979-2013.

For future climate data, we downloaded projections for 10 General Circulation Models (GCMs): (1) bcc-csm1-1, China; (2) CCSM4, USA; (3) CESM1-CAM5, USA; (4) CSIRO-Mk3-6-0, Australia; (5) GFDL-CM3, USA; (6) HadGEM2-AO, Korea; (7) IPSL-CM5A-MR, France; (8) MIROC-ESM-CHEM, Japan; (9) MIROC5, Japan; and (10) NorESM1-M, Norway (**Table S3**). We used median values across all 10 GCMs for all our analyses. By selecting multiple GCMs, we aimed to capture the uncertainty and variability around future climate scenarios. We selected two time periods 2050 (average for 2041-2060) and 2070 (average for 2061-2080), and two Representative Concentration Pathways (RCP) 4.5 and 6.0, which project a peak in emissions around 2040 and 2080, respectively, followed by a decline^51^. Of all GCMs, CSIRO-Mk3-6-0 showed the greatest variability for AP and PDQ (**Figure S15**).

### Species’ climate niche and cities’ climate

For the 16,006 species identified as being planted in urban areas, we used all occurrence records (i.e. within and outside urban areas) to extract values of the aforementioned climate variables to characterise species’ realised climate niches under baseline climatic conditions (i.e. 1979-2013). For each city, we placed a grid (1 × 1 km) over its area and extracted the values of all four variables at each cell for both current and future climates.

Then, for all species and cities, we estimated the upper and lower limits of the temperature and precipitation variables, respectively, based on all occurrence records (i.e. within and outside urban areas) for each species and based on all grid cells of each city. We used the upper and lower bounds of the distribution of values across the species range to determine whether cities are likely to exceed species’ limits. For this, we selected the threshold of the 95^th^ percentile of MAT/MTWM and the 5^th^ percentile of AP/PDQ. We used these thresholds to assess the extremes of these variables as indicative of niche tolerance (i.e. species’ thermal and drought stress tolerance for survival and growth)^4^. All throughout the manuscript, when referring to these climate variables, we imply the use of the 95^th^ (MAT/MTWM) and 5^th^ (AP/PDQ) percentiles, accordingly (**Figure S3**).

### Climate change vulnerability metrics

We selected three climate change vulnerability metrics for our analysis: exposure, safety margin and risk ^6,52^. These metrics were calculated for all four climate variables, time periods (baseline and future [2050 and 2070]) and RCPs (4.5 and 6.0).

Exposure (*E*) is a measure of how much the climate is projected to change (e.g. warmer or drier) between current and future climatic conditions; thus, it is estimated as the difference between the city’s future and baseline (i.e. current) climate as follow:

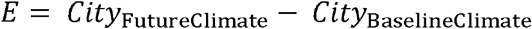

A positive exposure (*E* > 0 indicates that warmer (or wetter conditions are expected under future climate change scenarios.

The safety margin (*S* indicates how much warmer (or drier, a city could become before the realised climate niche of its species have been exceeded, and was estimated as follows:

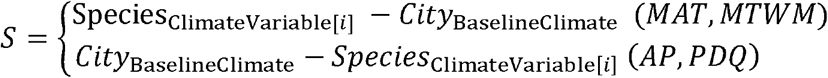

For S, a species’ climatic limit (Species_ClimateVariable[i]_ was measured as the 95^th^ (MAT/MTWM and the 5^th^ (AP/PDQ percentiles of the species’ climate niche based on its global occurrence records and baseline climatic conditions. The difference between Species_ClimateVariable[i]_ and the long-term average climatic conditions experienced in the focal city (i.e. City_BaselineClimate_ is calculated as the ‘safety margin’ (*S* for each focal species-by-city combination^4^. That is, a positive safety margin (*S* > 0 indicates that the species has a thermal tolerance limit which exceeds current baseline temperature conditions in the focal city (e.g. cooler and thus safe; whereas a negative value (*S* < 0 indicates that the species is experiencing “unsafe” climatic conditions under the baseline (e.g. warmer than what the species can actually withstand according to its tolerance limit for temperature (**Figure S3**.

The risk (*R* is the difference between *E* and *S*. Thus, if *R* is positive (*R* > 0, the exposure to future climate is greater than the current safety margin for the focal species in a focal city (i.e. high risk. Yet, if the difference is negative (*R* < 0 then exposure (*E* to future climate change is still within the range of values allowed by the safety margin (*S*), thus it is “safe” under future conditions (i.e. low risk (**Figure S3**. Risk to climate change (*R* was estimated as:

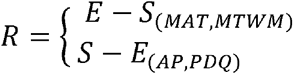

Linear regressions were fitted to evaluate the relationship between climate exposure/risk and cities’ latitude. Model performance was evaluated through the calculation of an R^2^ value and the *F*-Statistic at a significance level of *P* < 0.05. All analyses were conducted using the statistical software R version 4.0.5^45^. Caveats and limitations to our methodology can be found in the **Supplemental Materials** and prior studies^4,6^.

## Data availability

Global occurrence records are available on Global Biodiversity Information Facility (GBIF, www.gbif.org). Polygons defining the boundaries of 6,018 urban areas (are available on World Urban Areas (https://earthworks.stanford.edu/catalog/stanford-yk247bg4748). Climate data are available on CHELSA (https://chelsa-climate.org).

## Competing Interests

The authors declare that they have no conflict of interest to disclose.

## Author contribution

MER, RVG, PDR, SAP and MGT conceived the article. MER, RVG, LJB, PDR, SAP and MGT designed the research. MER, JBB, JL and BR collected and analysed data. MER wrote the article. All authors contributed to discussion of the content and reviewed or edited the manuscript before submission. All authors, except for MER and RVG, are listed alphabetically.

## Supplemental Material

### Results

#### Exposure

For MTWM, an average increase of 1.1°C (± 0.5°C) is predicted across all cities, rising to 1.9°C (± 0.6°C) by 2070 (**Table S1**). By 2050, the warmest months in four cities (Kiev, Ukraine; Belgorod and Bryansk, Russia; Belgrade, Serbia) are predicted to be >3°C warmer than baseline conditions (**Figure 1**). By 2070, 384 cities are projected to become >3°C warmer, with 13 predicted to be >4°C warmer than the baseline (average 1979-2013). For PDQ, 531 cities are predicted to become drier by 2050 (−4.2 mm ± 6.7 mm), and 531 cities will remain drier by 2070 (−5.8 mm ± 9.3 mm) (**Figure 2**).

**Figure 1.**
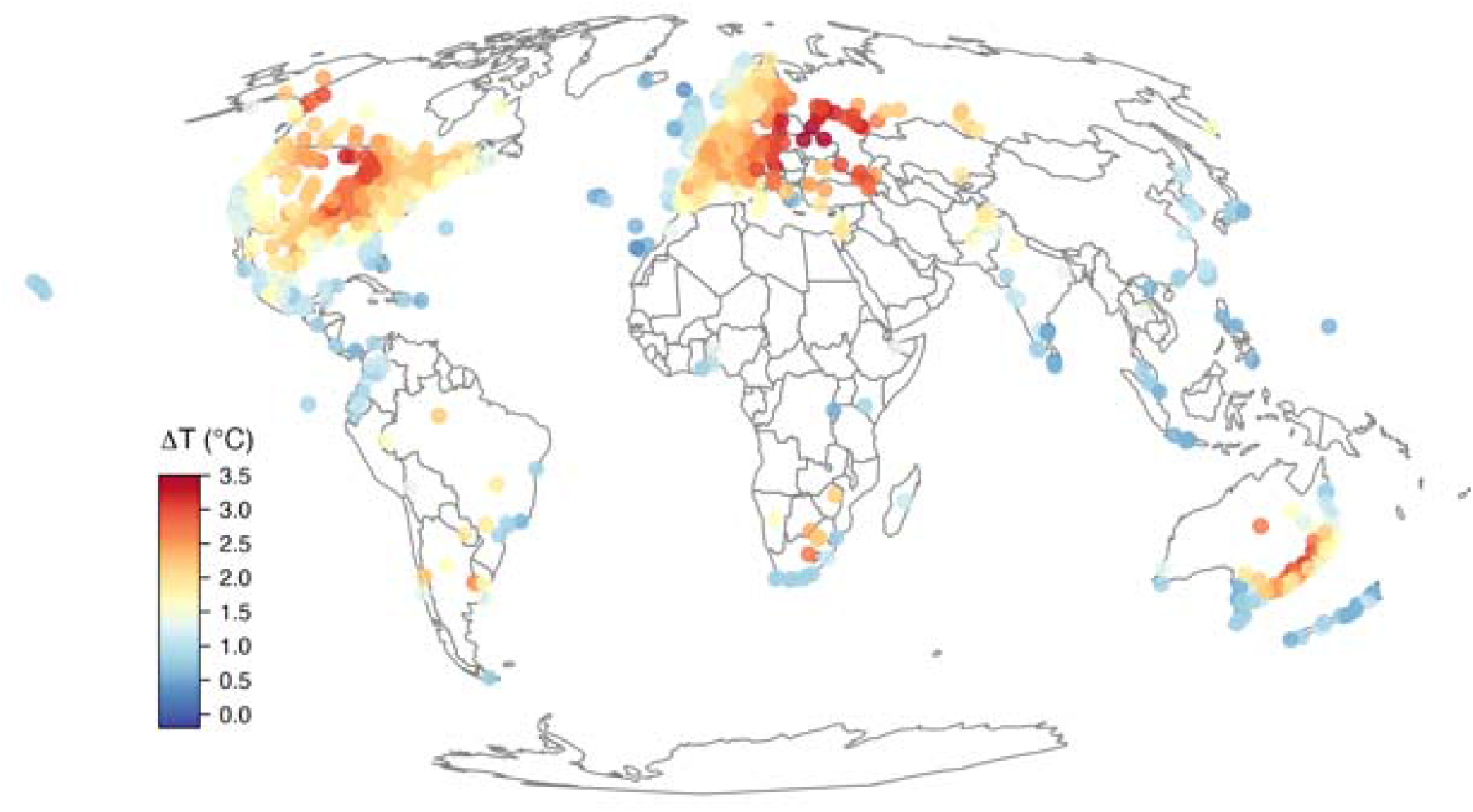
Changes (i.e. exposure) in maximum temperature of the warmest month (MTWM) predicted to occur by 2050. Data for RCP6.0.

**Figure 2.**
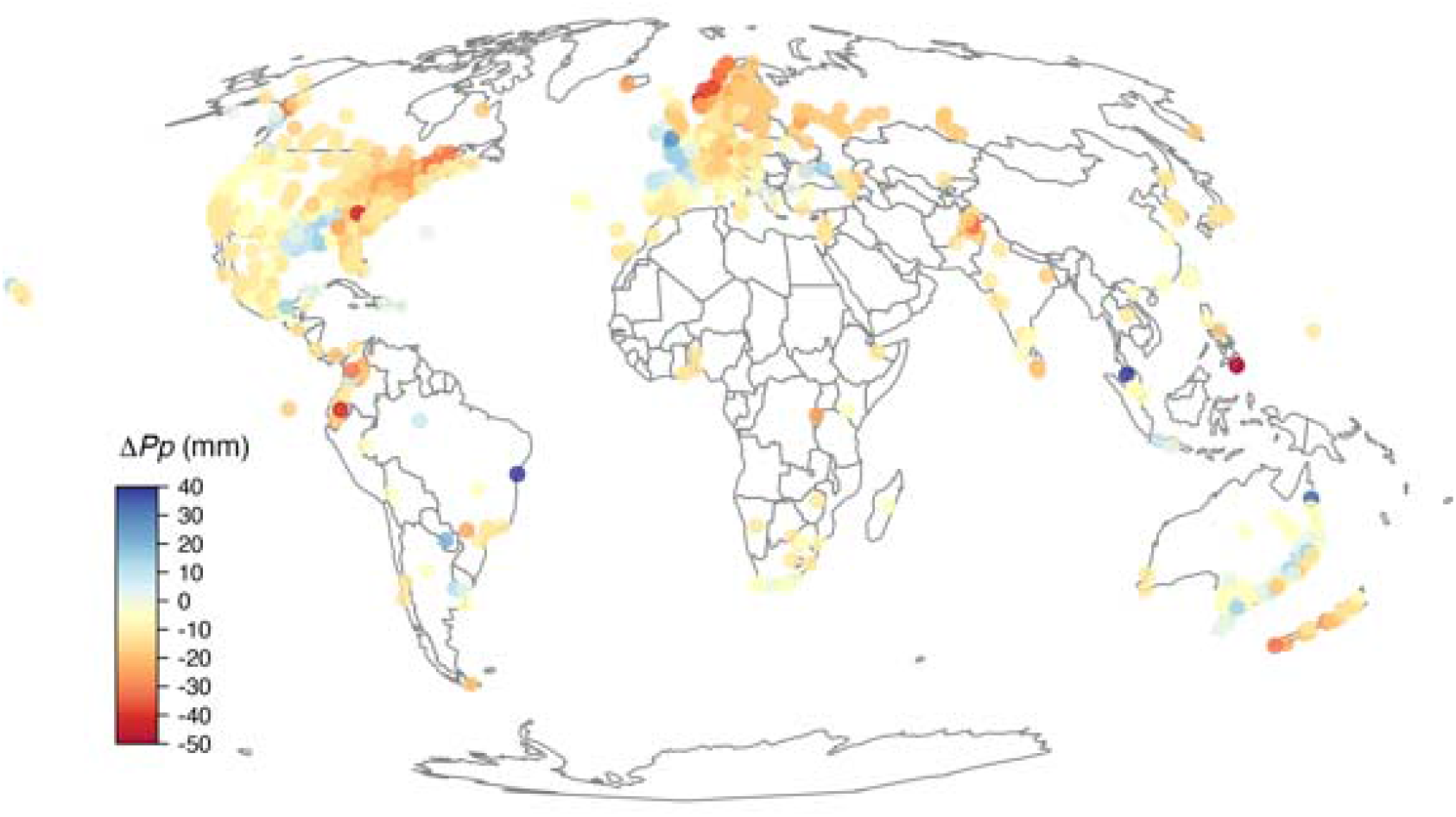
Changes (i.e. exposure) in precipitation of the driest quarter (PDQ) predicted to occur by 2050. Data for RCP6.0.

#### Safety margin

For MTWM, 9,005 species (56.3%) are exceeding their current MTWM safety margin. One city, Arrecife (Spain), has 100% of species exceeding their safety margin, and 122 cities have >50% of their species exceeding their safety margin (**Figure 3**). For PDQ, 7,487 species (46.8%) are exceeding their PDQ safety margin, in at least one city where they are currently planted, and 123 cities currently have >50% of their species exceeding their PDQ safety margin (**Figure 4**).

**Figure 3.**
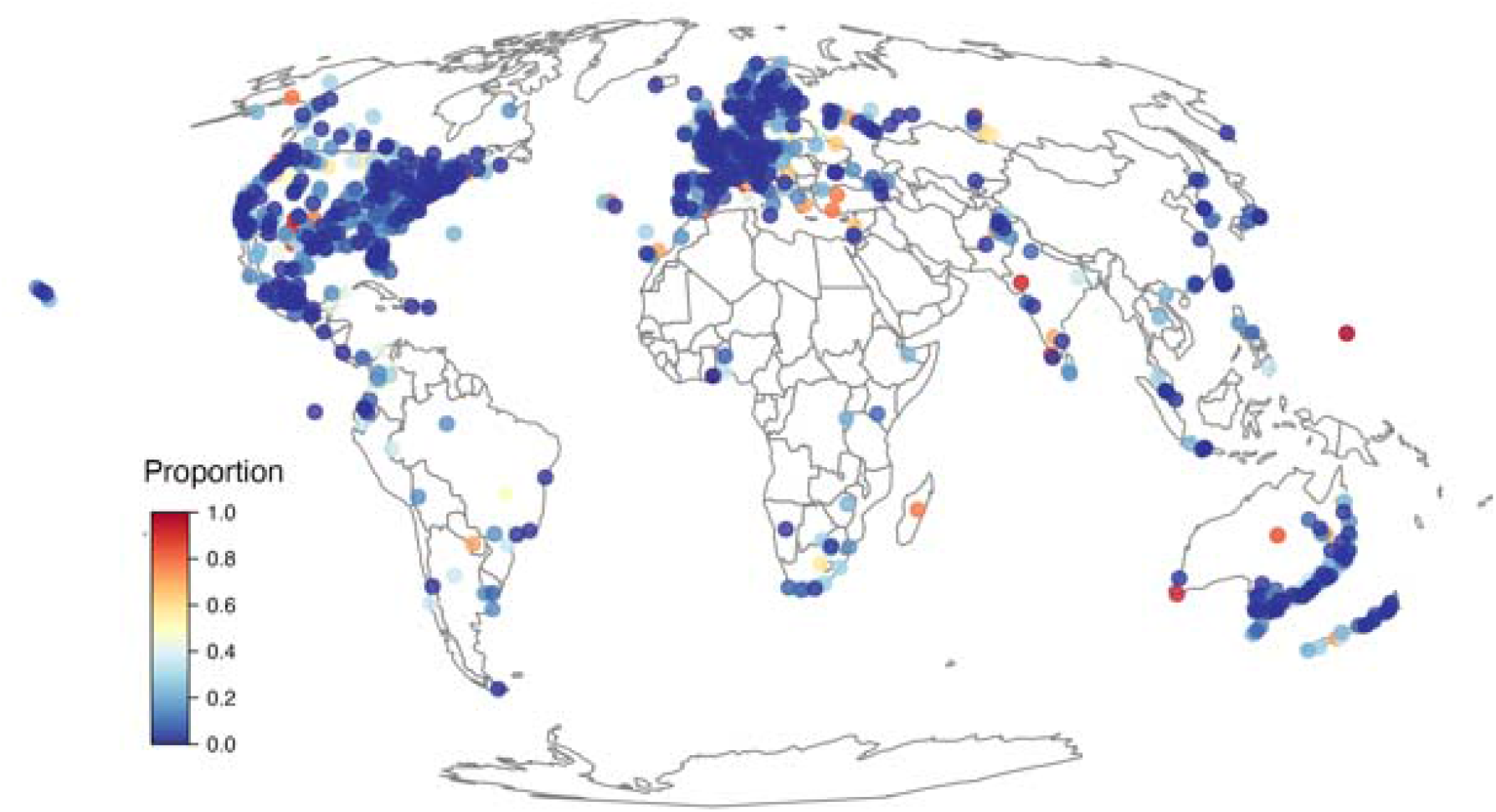
Proportion of urban forest species in each city predicted **to exceed their maximum temperature of the warmest month (MTWM) safety margin. Data for 2050 and RCP6.0.**

**Figure 4.**
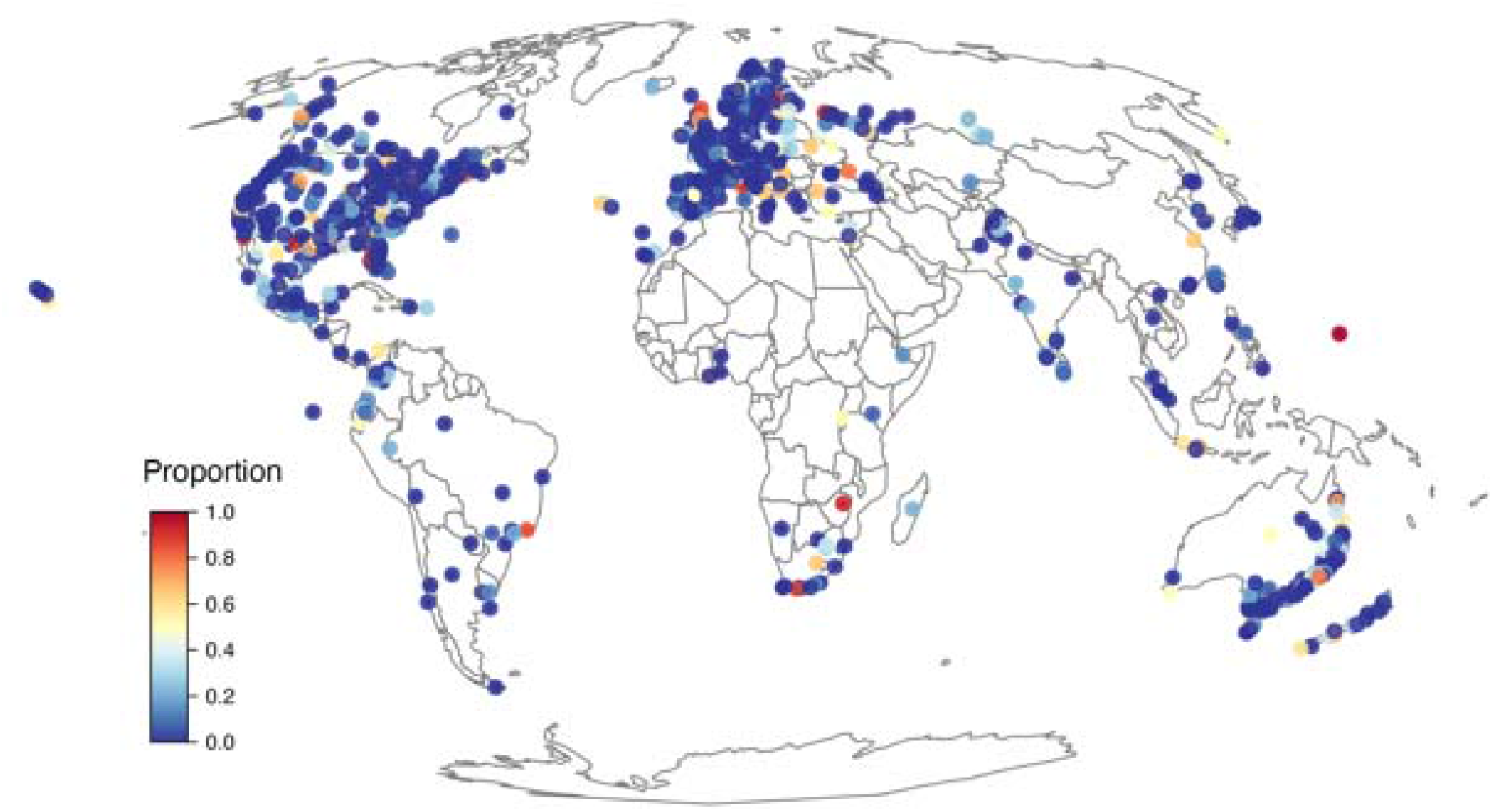
Proportion of urban forest species in each city predicted **to exceed their precipitation of the driest quarter (PDQ) safety margin. Data for 2050 and RCP6.0.**

#### Risk

By 2050, for MTWM and PDQ, 12,653 (79.1%) and 8,131 (50.8%) species, respectively, are predicted to be at risk from future changes in climate. For MTWM, 14 and 480 cities are predicted to have 100% and >50% of their species at risk. Whereas for PDQ, 10 and 131 cities are predicted to have 100% and >50% of their species at risk, respectively (**Figures 5–6**).

**Figure 5.**
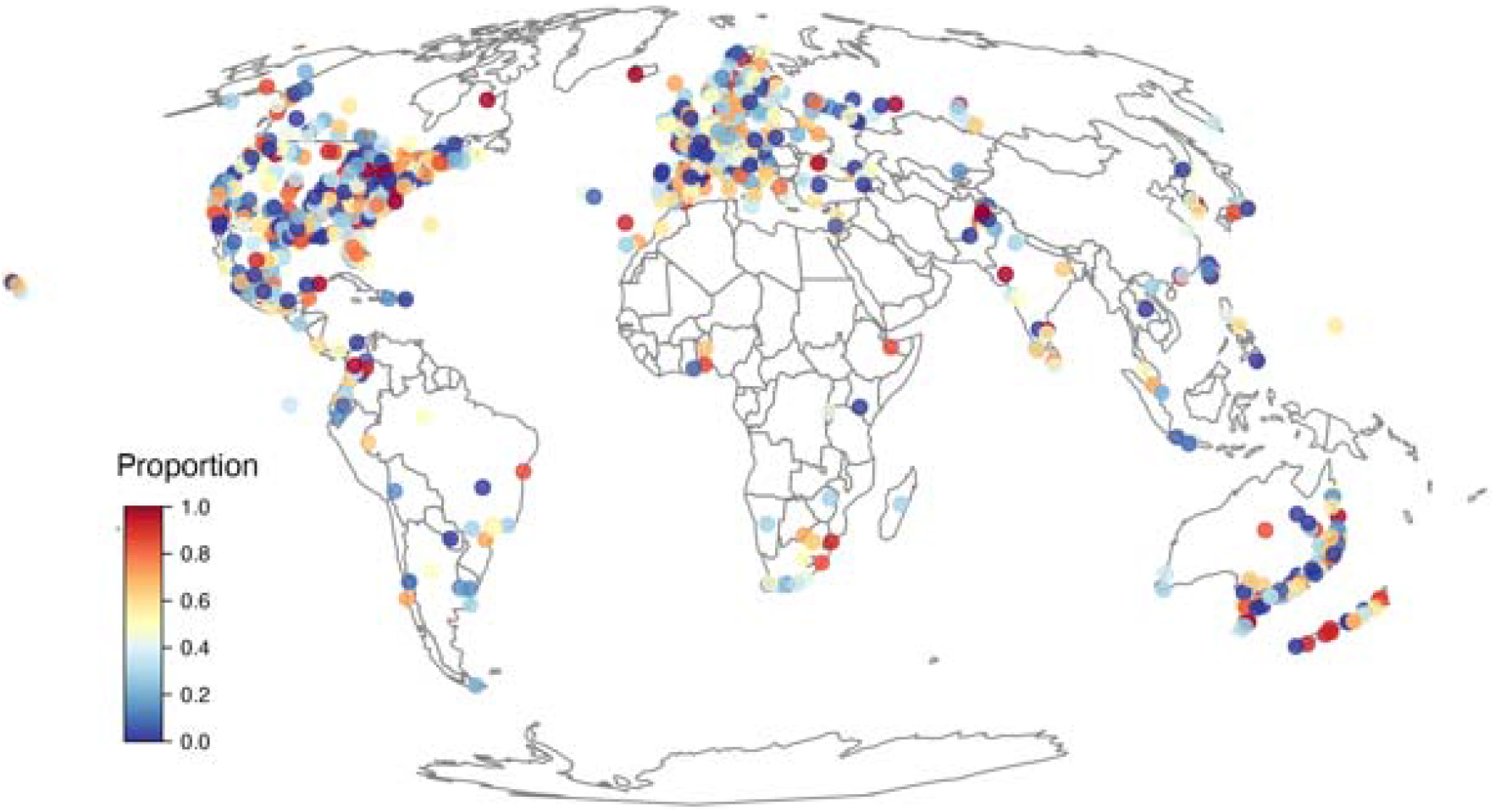
Proportion of urban forest species in each city predicted to be at risk from projected changes in **maximum temperature of the warmest month (MTWM). Data for 2050 and RCP6.0.**

**Figure 6.**
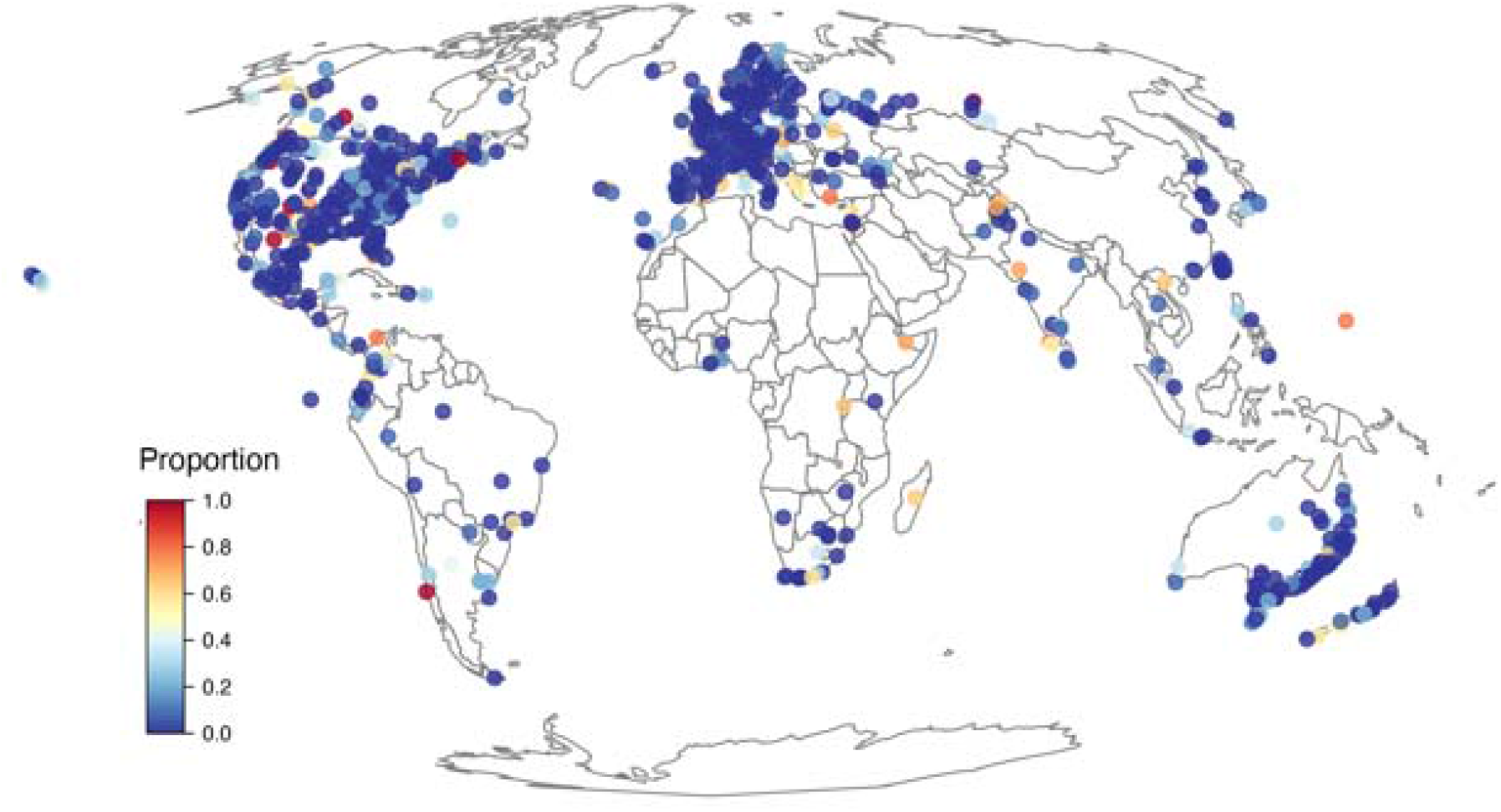
Proportion of urban forest species in each city predicted to be at risk from projected changes in **driest quarter (PDQ). Data for 2050 and RCP6.0.**

#### Caveats

##### Sampling bias

Sampling effort in herbarium collections largely reflects spatial variation, often related to human settlement and infrastructure^1,2^; hence, the sampling effort across cities is undoubtedly biased. Therefore, we acknowledge that our approach underestimates the number of species occurring within each city.

To assess the effect of potentially inadequate sampling on our analyses, we estimated the completeness of the species inventory in each city. Inventory completeness was calculated using an estimator of sample coverage^3^, which can be interpreted as the probability that a new occurrence record would not result in the observation of a previously unrecorded species. We then assessed the relationship between inventory completeness, and mean temperature and precipitation throughout urban areas (i.e. cities). For this assessment, temperature and precipitation variables were averaged for each city (i.e. averaging across all species in a given city) and plotted against inventory completeness aiming to identify a trend in variability in climate means as completeness increased. A variable that is sensitive to inventory completeness would show a “funnel effect” with higher variability at low completeness, thereby indicating biased samples (**Figure 7**).

**Figure 7.**
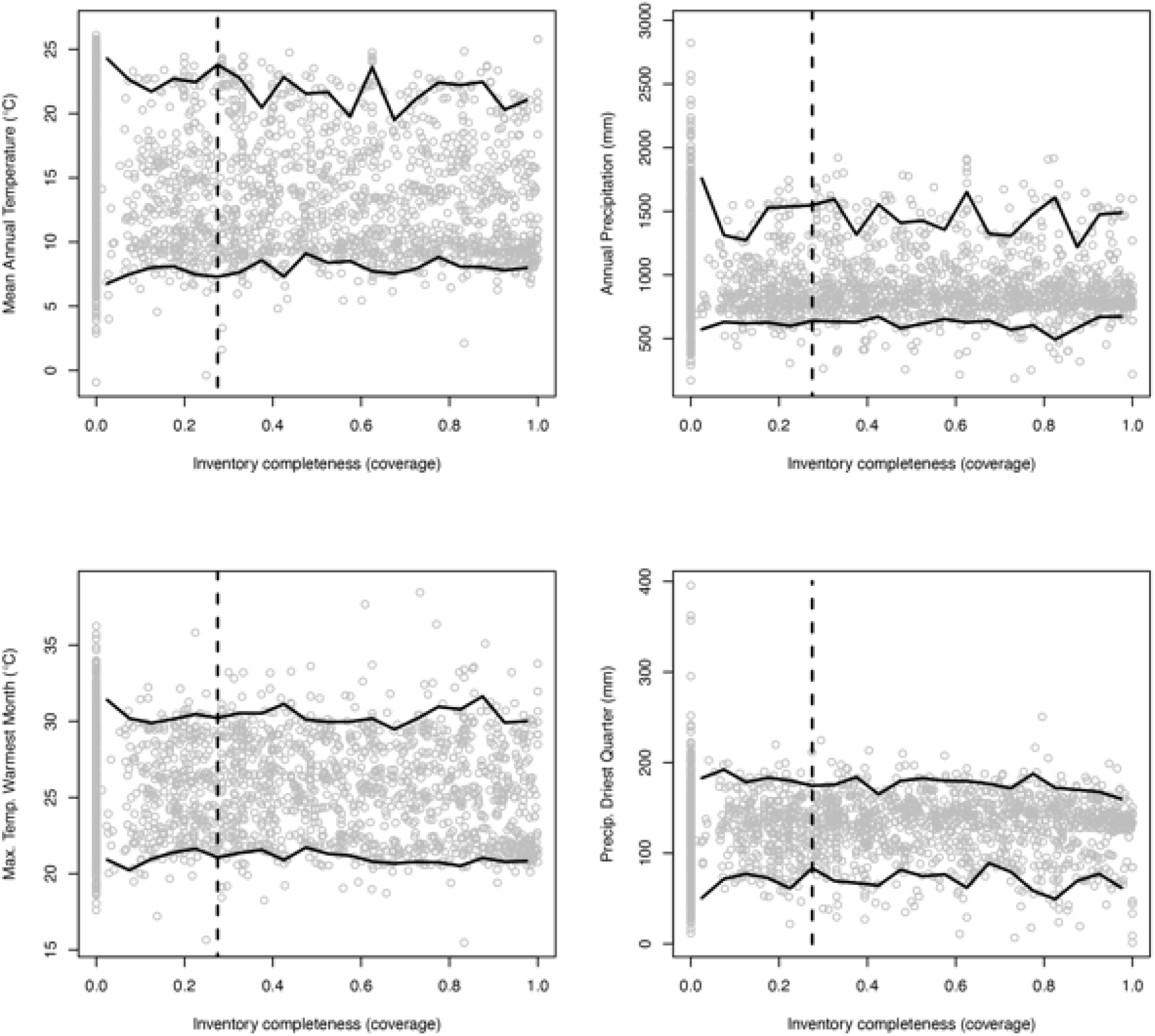
Average climate value for each city (mean annual temperature, annual precipitation, maximum temperature of the warmest month, and precipitation of the driest quarter) plotted against the cities’ species coverage. For each plot, we plotted the 5% and 95% quantiles for coverage bins (i.e. 0-0.05, 0.05-0.10, 0.10-0.15, etc.) and the median coverage.

Although our results indicated that the climate variables are robust to inventory completeness, we excluded cities with low completeness (<0.20) from our analyses, retaining 1,010 cities from 93 countries. The average number of cities across countries was 11 (standard deviation ± 35 cities), with a maximum of 324 in the USA, while 44 countries were represented by a single city. The average human population across all 1,010 cites was 830,834 (± 2,193,136 people), with a maximum of 35,676,000 in Tokyo (Japan) and a minimum of 246 people in Theodore (Australia).

#### Methodological approach

First, species realised niches (assessed here) are not equal to their fundamental niche. We used occurrence records to approximate distribution, but biological factors such as competition, abiotic factors (e.g. soil and nutrients) and dispersal limits may mean that our use of occurrence records underestimate species’ climate envelopes. Second, climate data based on coarse-grained spatial interpolations from weather stations that are shielded from direct solar radiation used here, fail to identify areas where harsh conditions can be exacerbated due to the urban heat island effect^4^ or areas where conditions are more benign due to the vegetation effect on microclimate (e.g. high canopy cover)^5^ or presence of wind^6^, the latter is particularly important in coastal cities. Third, our assessment did not consider other environmental factors that can mitigate or exacerbate the effects of climate change, such as the presence/absence of water bodies, soil type and topography. Fourth, our approach does not consider species’ adaptive capacity, which facilitate species’ resilience to climate change, and the potential feedback mechanisms between climate and biota (e.g. the role of vegetation in modulating temperature). Additional factors that were not considered in our approach, such as urban heat island effect, sea level rise, human impacts (e.g. exploitation and pollution), land-use change and deforestation, also erode the resilience of cities to climate change. Finally, by selecting a more conservative scenario (i.e. RCP6.0) following Raftery, et al. ^7^, our predictions might underestimate the changes in climate, compared to a less conservative scenario, such as RCP8.5^8^.

## Supplemental Tables and Figures

**Supplemental Table S1.**
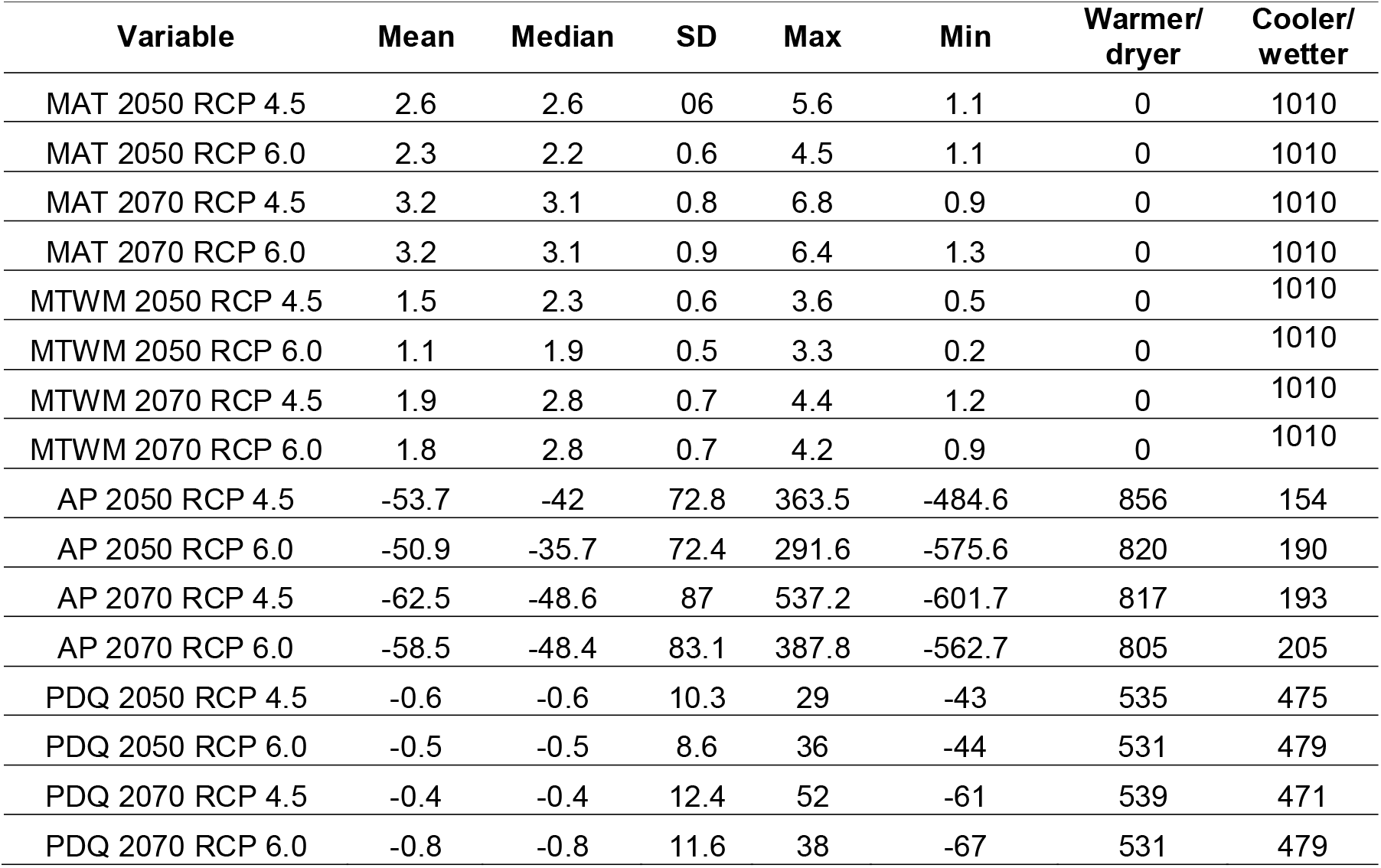
Summary of mean, median, standard deviation (SD) maximum (Max) and minimum (min) values estimated for two time periods (2050 and 2070) and two Representative Concentration Pathway (RCP 4.5 and 6.0) for mean annual temperature (MAT; °C), maximum temperature of the warmest month (MTWM; °C), annual precipitation (AP; mm), and precipitation of the driest quarter (PDQ; mm) across 1010 cities; and number of cities that are predicted to become warmer/dryer and cooler/wetter for each climate scenario.

| Variable | Mean | Median | SD | Max | Min | Warmer/<br>dryer | Cooler/<br>wetter |
| --- | --- | --- | --- | --- | --- | --- | --- |
| MAT 2050 RCP 4.5 | 2.6 | 2.6 | 06 | 5.6 | 1.1 | 0 | 1010 |
| MAT 2050 RCP 6.0 | 2.3 | 2.2 | 0.6 | 4.5 | 1.1 | 0 | 1010 |
| MAT 2070 RCP 4.5 | 3.2 | 3.1 | 0.8 | 6.8 | 0.9 | 0 | 1010 |
| MAT 2070 RCP 6.0 | 3.2 | 3.1 | 0.9 | 6.4 | 1.3 | 0 | 1010 |
| MTWM 2050 RCP 4.5 | 1.5 | 2.3 | 0.6 | 3.6 | 0.5 | 0 | 1010 |
| MTWM 2050 RCP 6.0 | 1.1 | 1.9 | 0.5 | 3.3 | 0.2 | 0 | 1010 |
| MTWM 2070 RCP 4.5 | 1.9 | 2.8 | 0.7 | 4.4 | 1.2 | 0 | 1010 |
| MTWM 2070 RCP 6.0 | 1.8 | 2.8 | 0.7 | 4.2 | 0.9 | 0 | 1010 |
| AP 2050 RCP 4.5 | -53.7 | -42 | 72.8 | 363.5 | -484.6 | 856 | 154 |
| AP 2050 RCP 6.0 | -50.9 | -35.7 | 72.4 | 291.6 | -575.6 | 820 | 190 |
| AP 2070 RCP 4.5 | -62.5 | -48.6 | 87 | 537.2 | -601.7 | 817 | 193 |
| AP 2070 RCP 6.0 | -58.5 | -48.4 | 83.1 | 387.8 | -562.7 | 805 | 205 |
| PDQ 2050 RCP 4.5 | -0.6 | -0.6 | 10.3 | 29 | -43 | 535 | 475 |
| PDQ 2050 RCP 6.0 | -0.5 | -0.5 | 8.6 | 36 | -44 | 531 | 479 |
| PDQ 2070 RCP 4.5 | -0.4 | -0.4 | 12.4 | 52 | -61 | 539 | 471 |
| PDQ 2070 RCP 6.0 | -0.8 | -0.8 | 11.6 | 38 | -67 | 531 | 479 |

**Supplemental Table S2.**
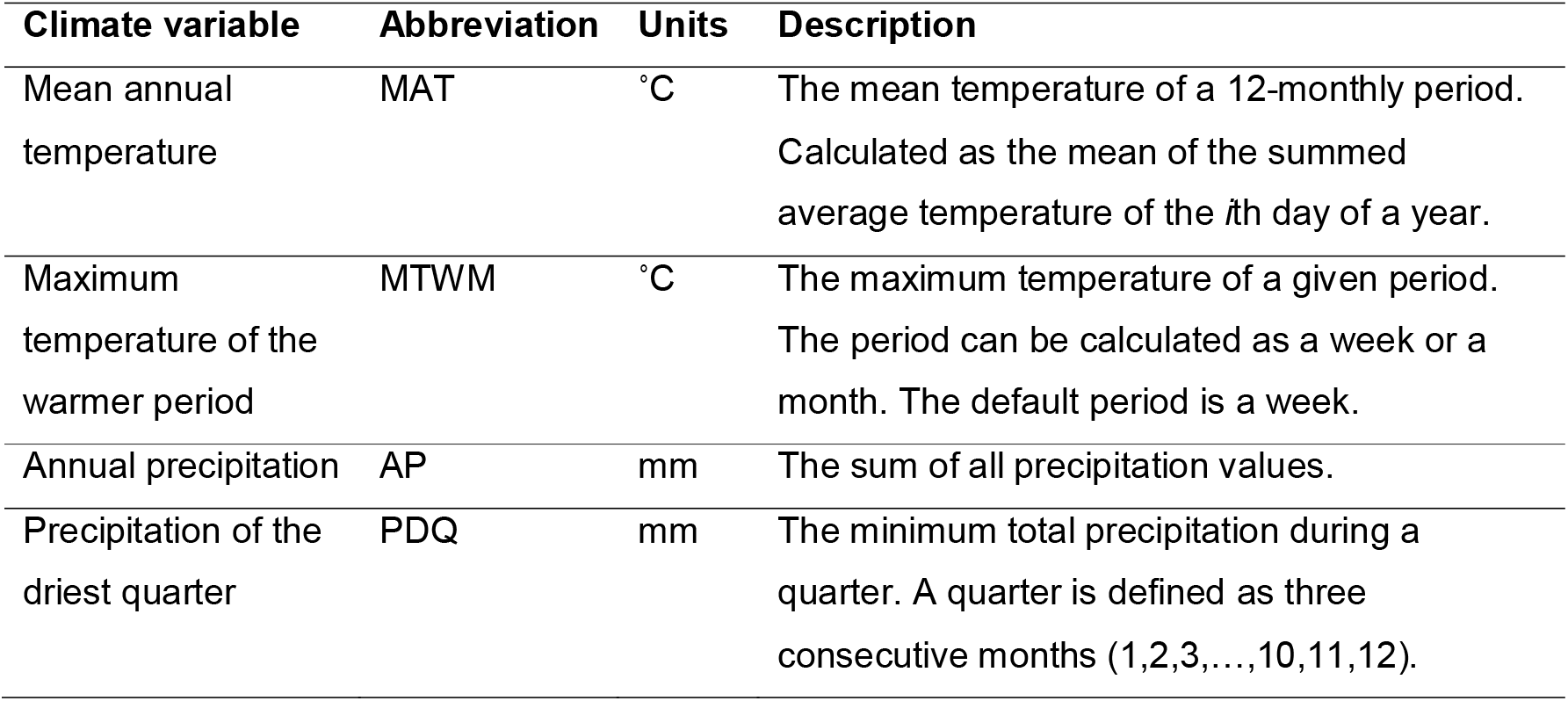
Climate variables selected for this study. More details on climate variables can be found in O’Donnell and Ignizio ^9^

| Climate variable | Abbreviation | Units | Description |
| --- | --- | --- | --- |
| Mean annual temperature | MAT | °C | The mean temperature of a 12-monthly period.<br>Calculated as the mean of the summed average temperature of the $i$th day of a year. |
| Maximum temperature of the warmer period | MTWM | °C | The maximum temperature of a given period.<br>The period can be calculated as a week or a month. The default period is a week. |
| Annual precipitation | AP | mm | The sum of all precipitation values. |
| Precipitation of the driest quarter | PDQ | mm | The minimum total precipitation during a quarter. A quarter is defined as three consecutive months (1,2,3,...,10,11,12). |

**Supplemental Table S3.** Global Circulation Models (GCMs) from the 10 CMIP5 models considered in this study.

| **GCM** | **Country** | **Reference** |
| --- | --- | --- |
| Bcc-csm1-1 | China | Wu, et al. <sup>10</sup> |
| CCSM4 | USA | https://www2.cesm.ucar.edu/models |
| CESM1-CAM5 | USA | https://www2.cesm.ucar.edu/models |
| CSIRO-Mk3-6-0 | Australia | Jeffrey, et al. <sup>11</sup> |
| GFDL-CM3 | USA | http://www.gfdl.noaa.gov/earthsystem-model |
| HADGEM2-AO | Korea | http://cms.ncas.ac.uk/wiki/UM/<br>Configurations/HadGEM2 |
| IPSL-CM5A-MR | France | http://icmc.ipsl.fr/index.php/<br>icmc-models/icmc-ipsl-cm5 |
| MIROC5 | Japan | Watanabe, et al. <sup>12</sup> |
| MIROC-ESM-CHEM | Japan | http://www.wcrp-climate.org/<br>wgcm/WGCM15/presentations/<br>21Oct/KIMOTO_Japan.pdf |
| NorESM1-M | Norway | https://verc.enes.org/ISENES2/<br>models/earthsystem-models/<br>ncc/noresm |

**Supplemental Figure S1.**
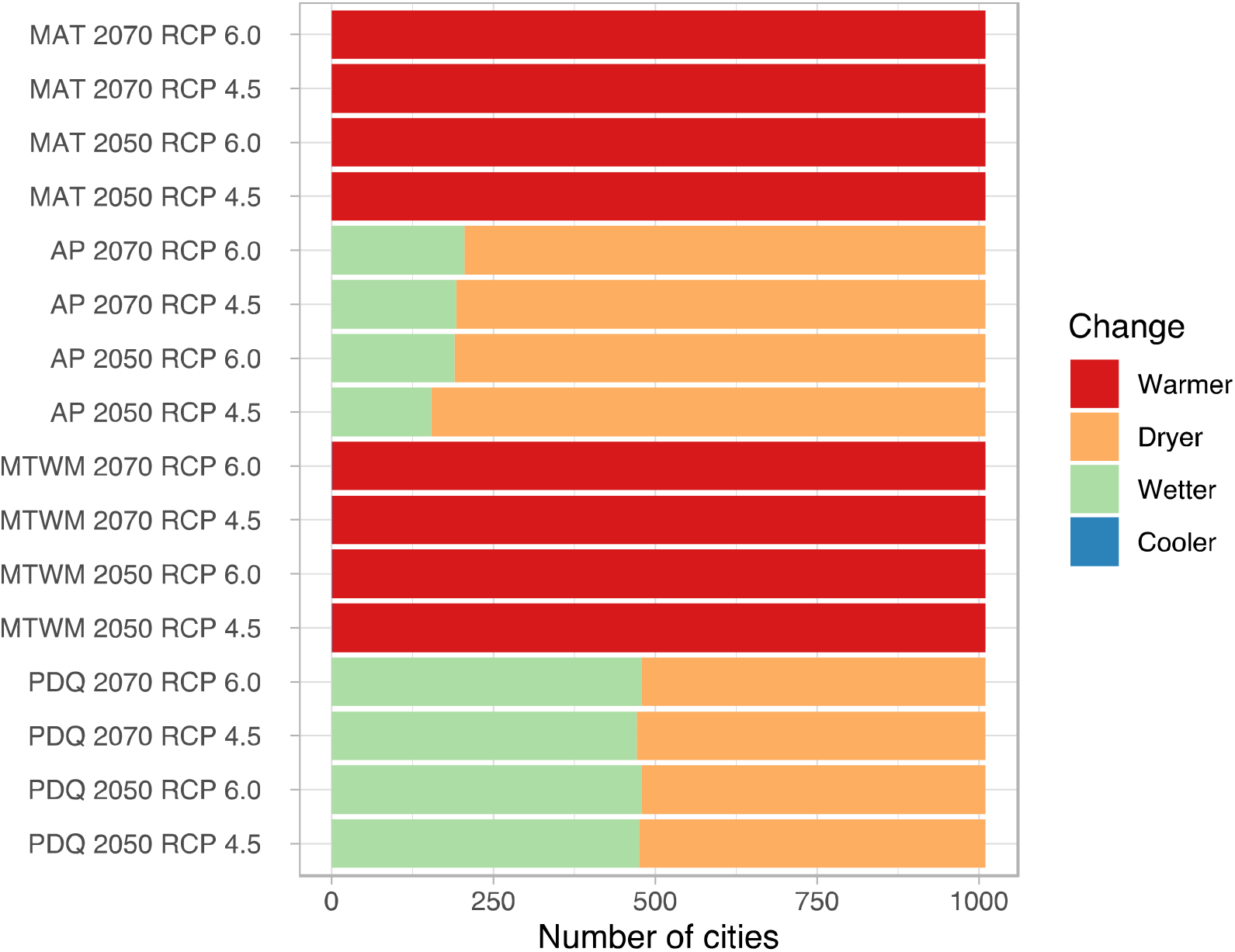
Number of cities (n= 1,010) predicted to become warmer/cooler and dryer/wetter in two time periods (2050 and 2070) and under two Representative Concentration Pathway (RCPs; 4.6 and 6.0). MAT = mean annual temperature, AP = annual precipitation, MTWM = maximum temperature of the warmest month, and PDQ = precipitation of the driest quarter.

**Supplemental Figure S2.**
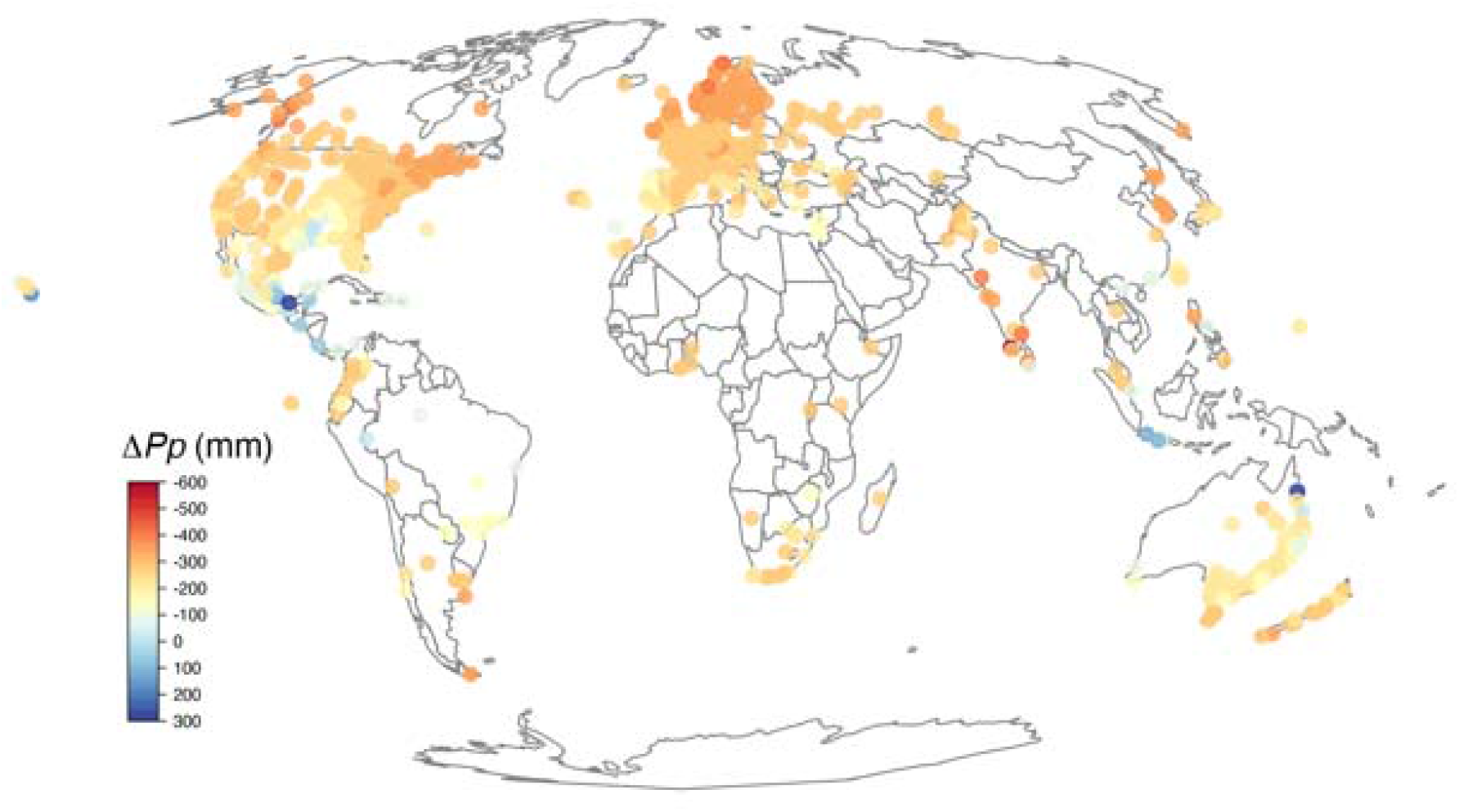
Changes (i.e. exposure) in annual precipitation (AP) predicted to occur by 2050. Data for RCP6.0.

**Supplemental Figure S3.**
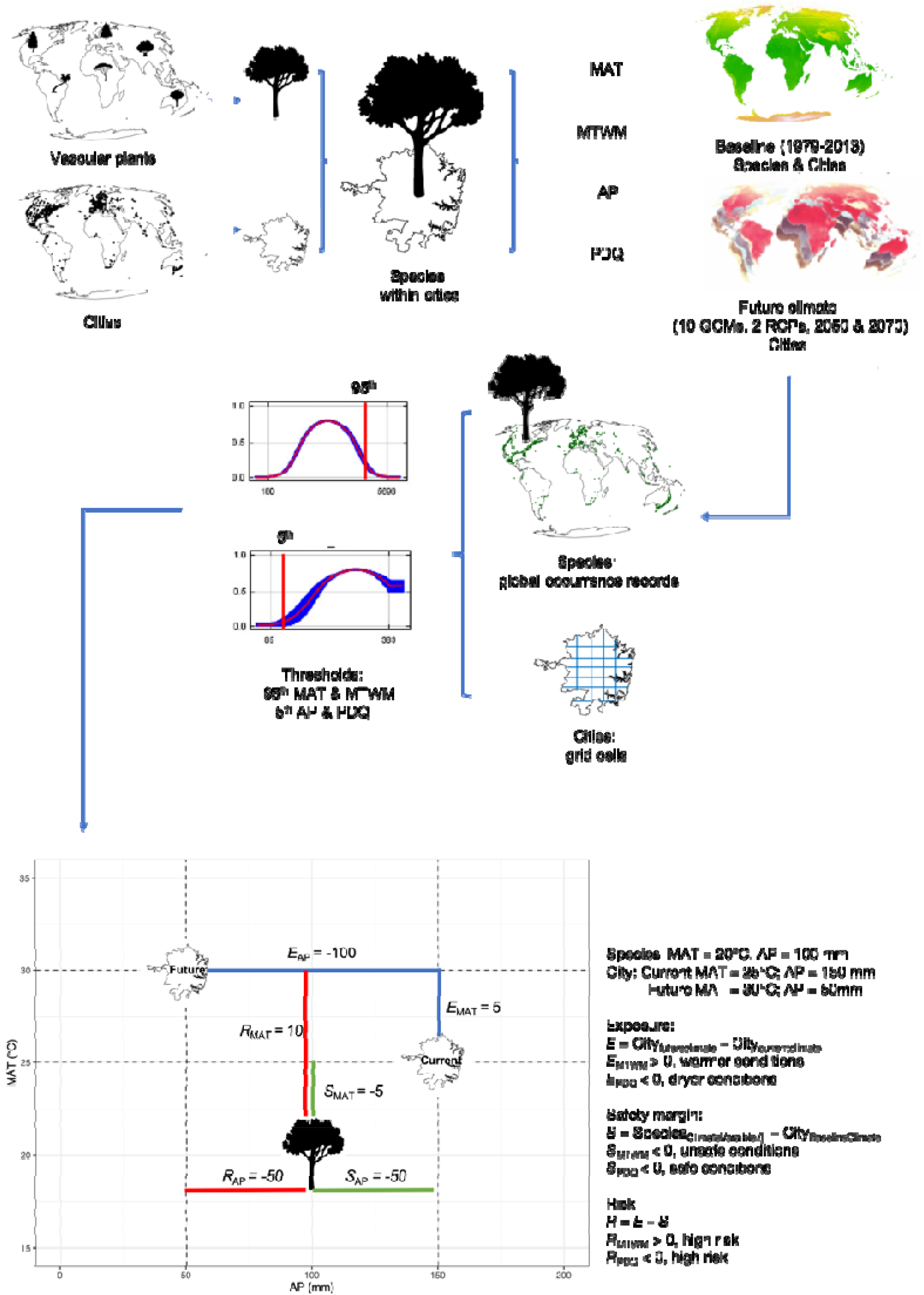
We obtained occurrence records for all tracheophytes and polygons defining the boundaries of cities globally. Then, we identified species occurring only within cities (i.e. 16,006 species). Climate data were downloaded for four climate variables (**mean annual temperature [MAT; °C], maximum temperature of the warmest month [MTWM; °C], annual precipitation [AP; mm], and precipitation of the driest quarter [PDQ; mm]) for baseline (average for** 1979-2013) and future conditions (10 **General Circulation Models** [GCMs], two time periods 2050 [average for 2041-2060] and 2070 [average for 2061-2080], and two Representative Concentration Pathways [RCP] 4.5 and 6.0). We used baseline climate to estimate species’ realized niches from all global occurrence records (i.e. within and outside cities). Baseline and future climate were used to estimate cities’ climate from a grid (1 × 1 km) placed over its area. We then estimated the threshold of the 95^th^ percentile of MAT/MTWM and the 5^th^ percentile of AP/PDQ to assess the extremes of these variables as indicative of niche tolerance. For each city, we estimated its exposure to climate change (i.e. a measure of how much the climate is projected to change [e.g. warmer and drier] between current and future climatic conditions); in this example, the city will become 5°C warmer (E_MAT_) and decrease 100 mm in rainfall (E_AP_). For each species at each city, we estimate its safety margin; i.e. an index of how much warmer (or drier) a city could become before the realized climate niches of its species is exceeded; in this example, the species is currently experiencing unsafe conditions for MAT, as the city threshold (City_Current95thMAT_ = 25°C) is 5°C warmer than the species’ threshold (Species_95thMAT_ = 20°C); whereas for precipitation, the species is experiencing safe conditions, as the species’ threshold (Species_5thAP_ = 100 mm) is lower than the city’s threshold (City_Current5thAP_ = 150 mm). Finally, risk was estimated as the difference between exposure and safety margin (*R* = *E* – S); in this example, the city is becoming warmer and dryer and for both variables the species will be at high risk. By 2050 the city will be 10°C warmer than the species MAT threshold and 50 mm dryer than the species AP threshold.

**Supplemental Figure S4.**
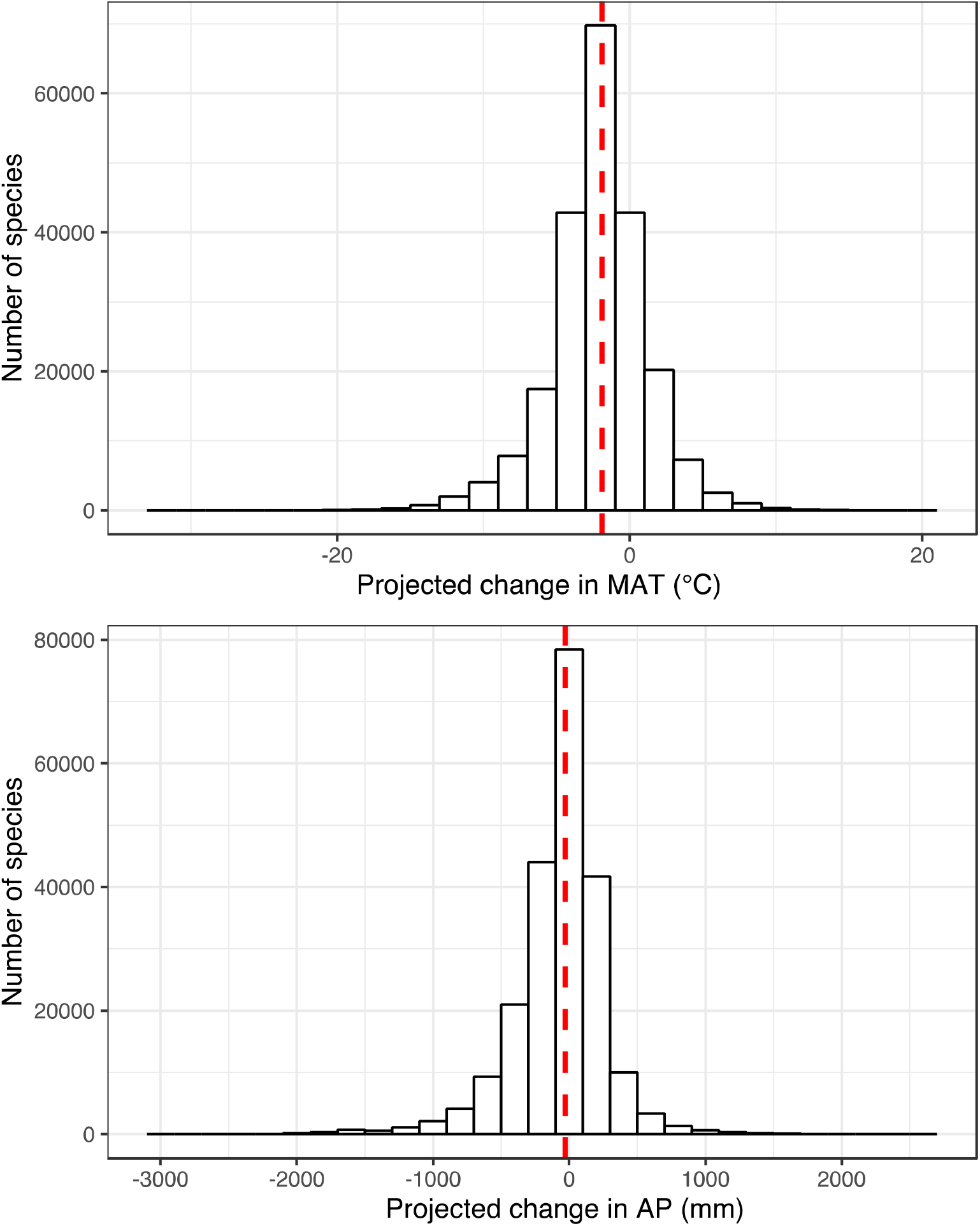
Changes in mean annual temperature (MAT) and annual precipitation (AP) projected to occur in species’ safety margin across 1,010 cities across the world by 2050 (RCP6.0). Note that the count of species exceeds the number of species, as one species can have a different safety margin depending on the city where it is planted. Median across all data is marked in red line.

**Figure S5.**
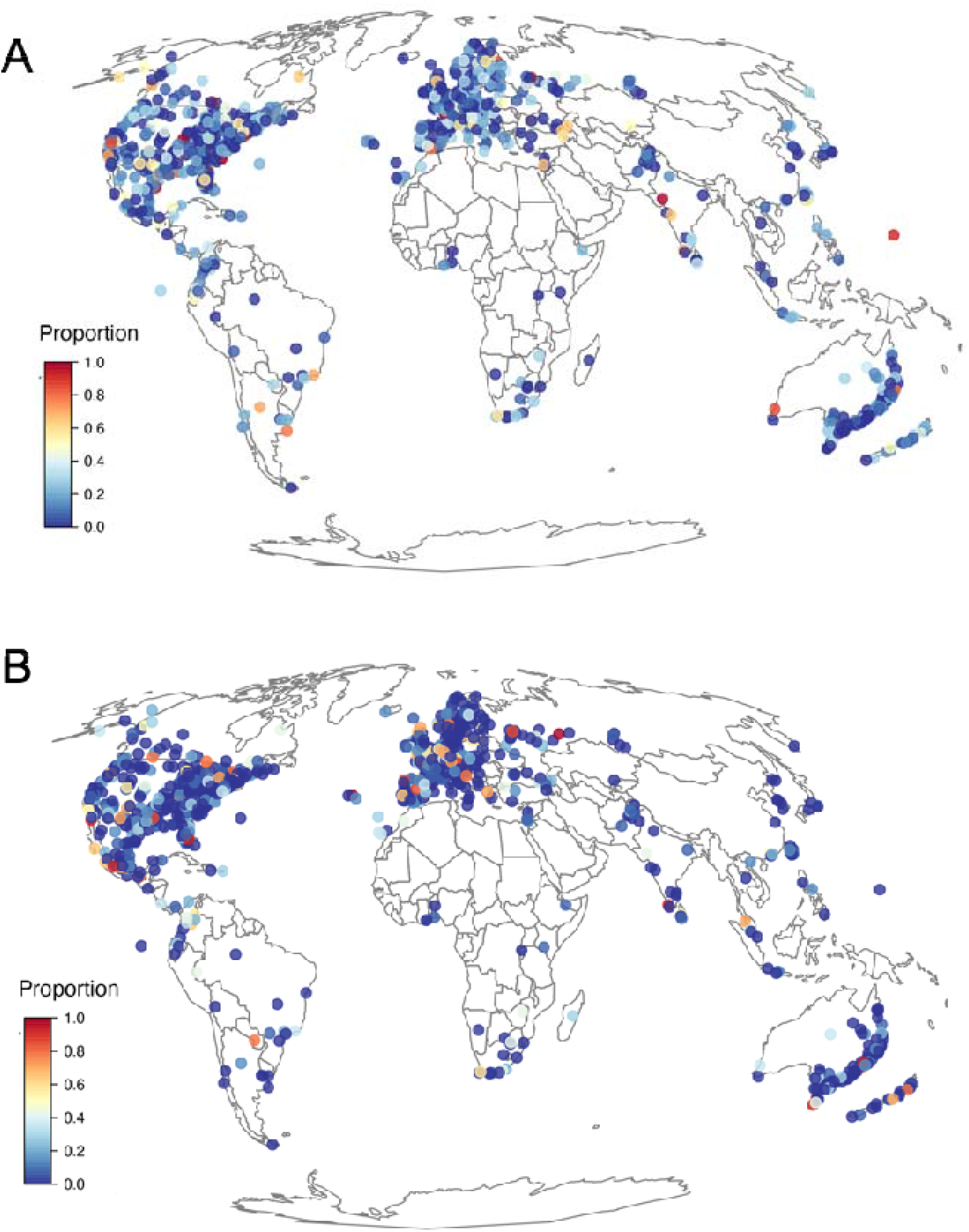
Proportion of urban forest species in each city predicted **to exceed their mean annual temperature (A) and annual precipitation (B) safety margin.**

**Supplemental Figure S6.**
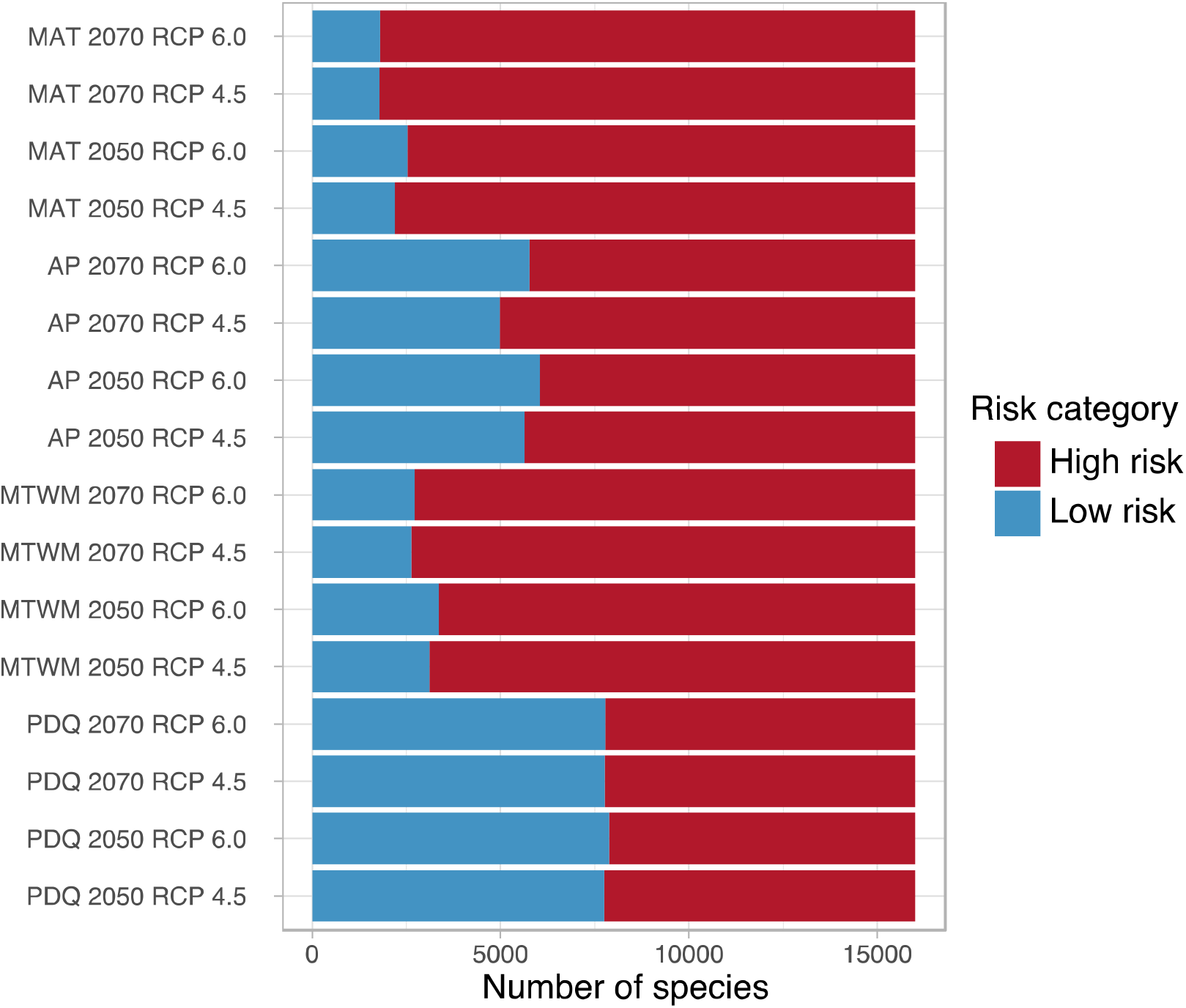
Number of species identified to be at high and low risk of climate change for four climate variables: mean annual temperature (MAT); annual precipitation (AP); maximum temperature of the warmest month (MTWM); and precipitation of the driest quarter (PDQ); two time periods (2050 and 2070) and two Representative Concentration Pathway (RCP 4.5 and 6.0).

**Supplemental Figure S7.**
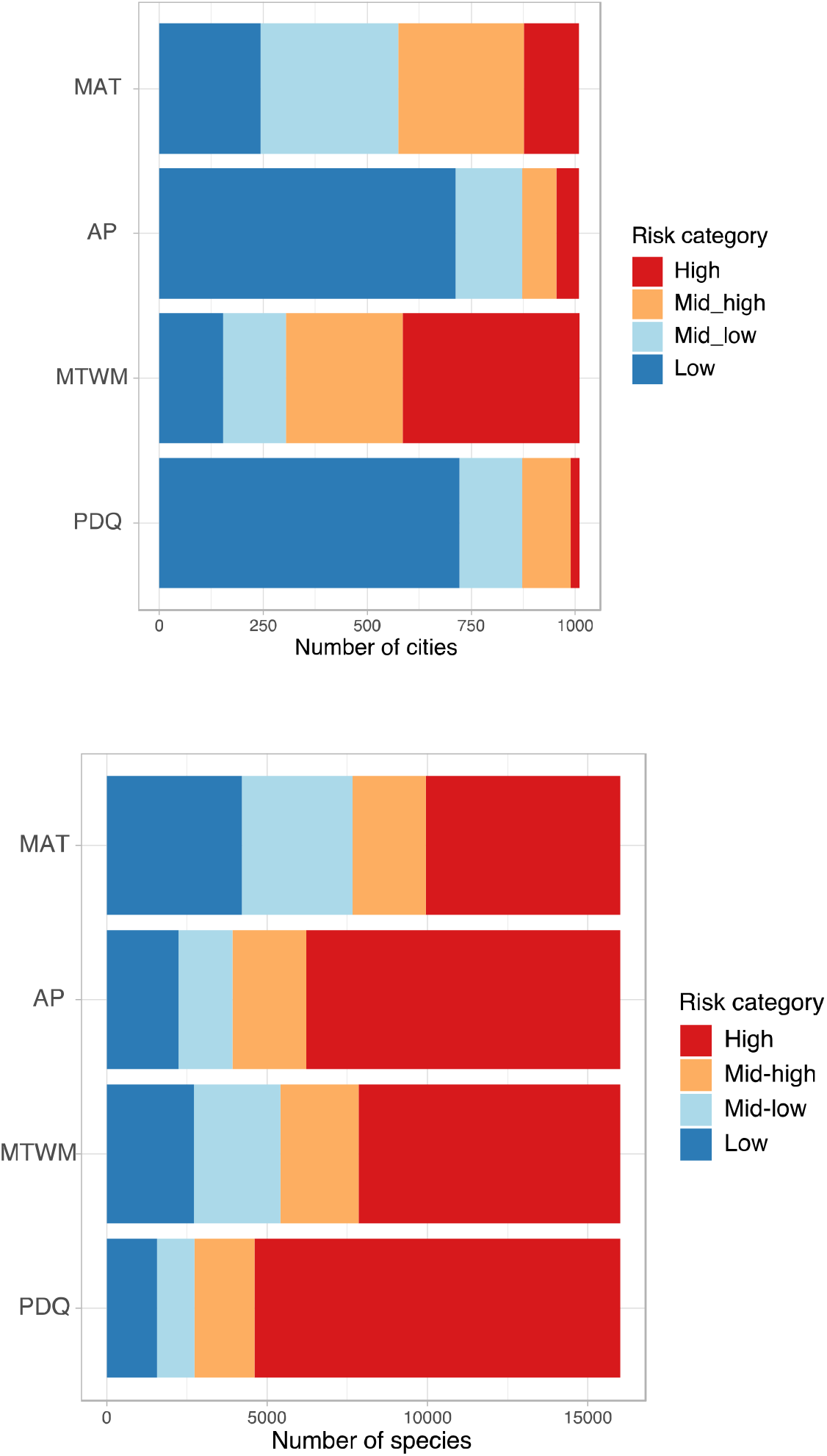
Number of cities with proportion of species in four risk categories: high (>75%), mid-high (75-50%), mid-low (50-25%) and low (<25%) (A); and number of species with proportion of cities at each risk category (B). Data for 2050 and RCP6.0.

**Supplemental Figure S8.**
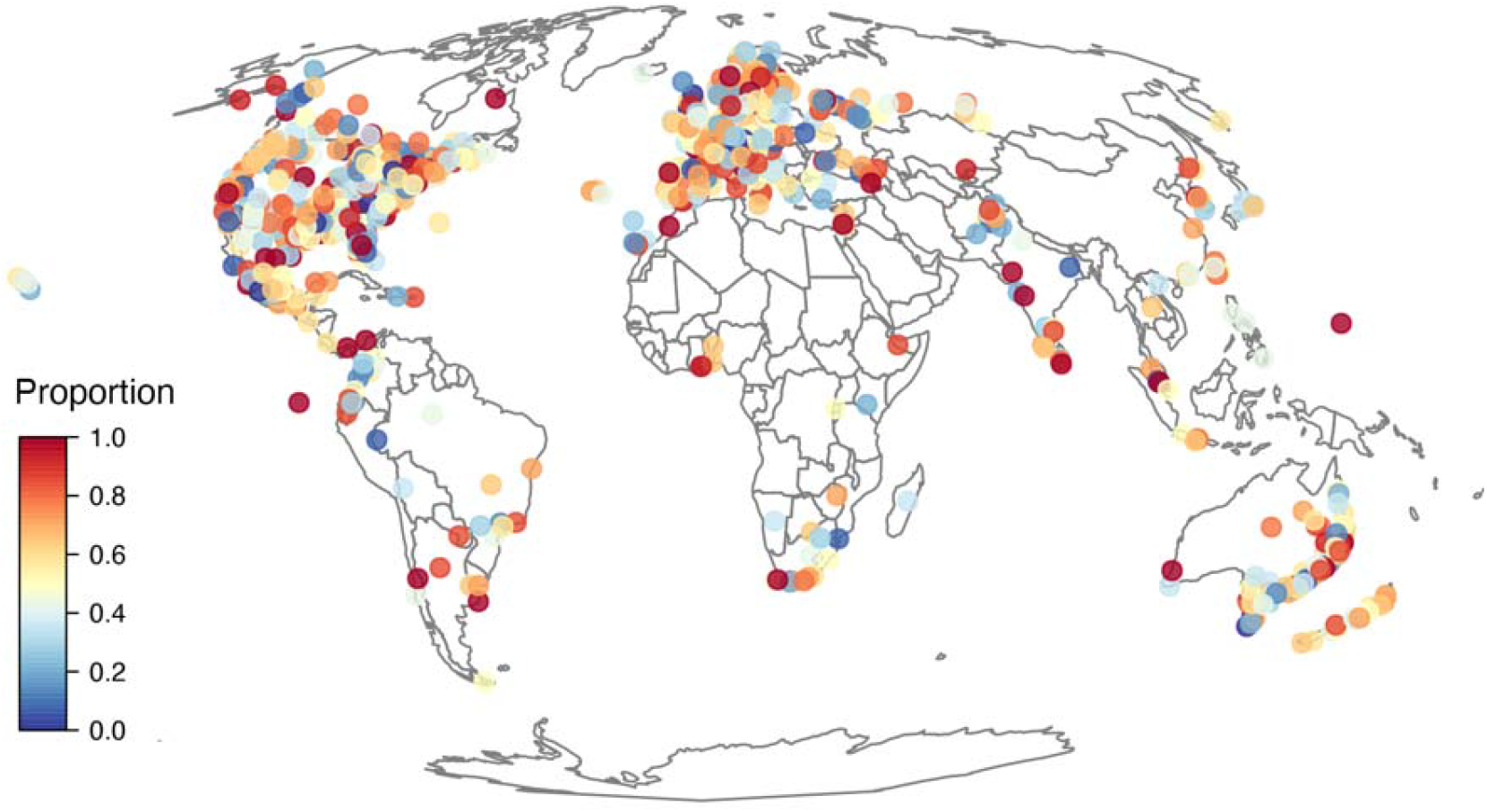
Proportion of urban forest species in each city predicted **to be at risk of future change in mean annual temperature (MAT) by 2070. Data for RCP6.0.**

**Supplemental Figure S9.**
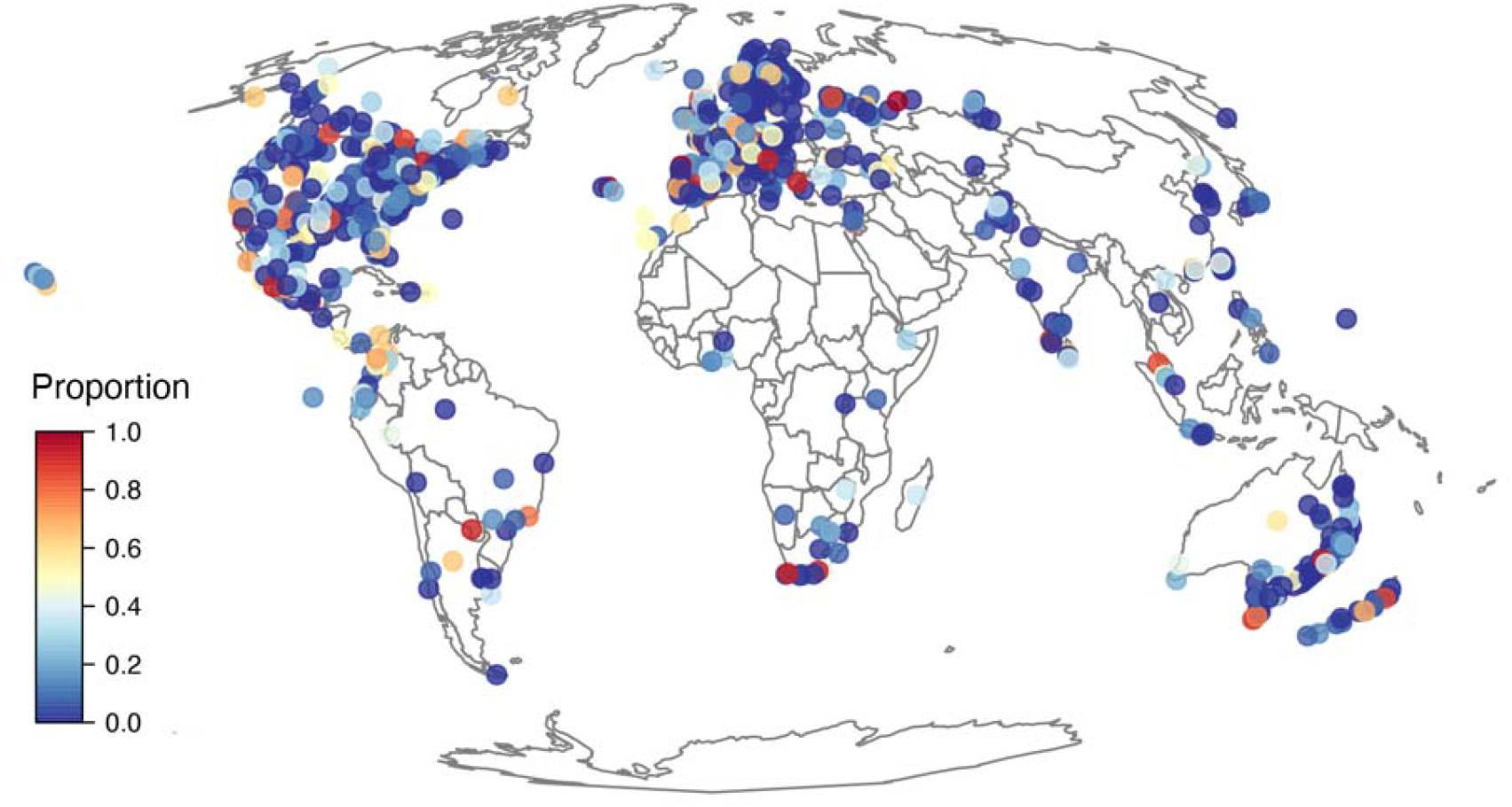
Proportion of urban forest species in each city predicted **to be at risk of future change in annual precipitation (AP) by 2070. Data for RCP6.0.**

**Supplemental Figure S10.**
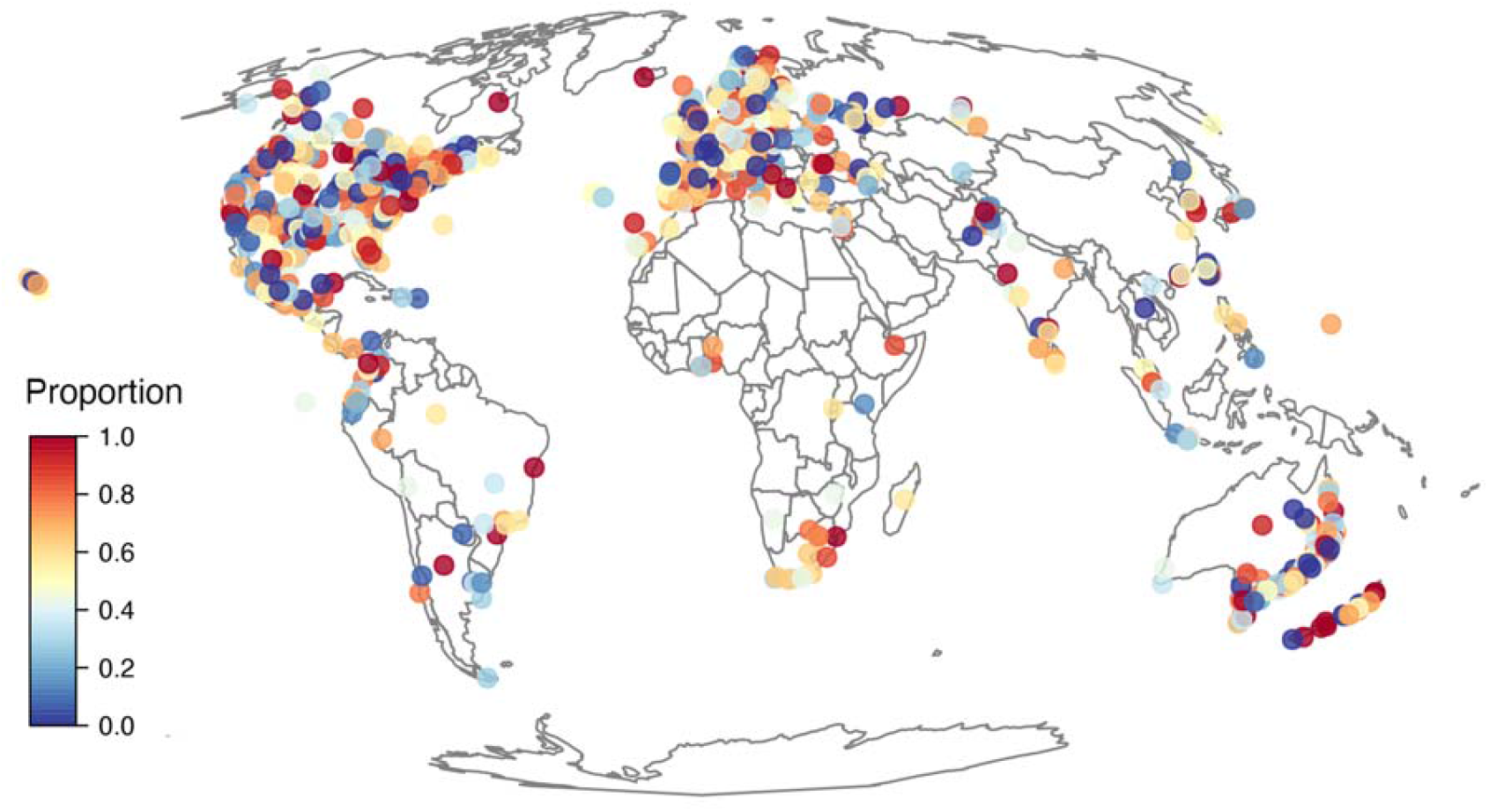
Proportion of urban forest species in each city predicted **to be at risk of future change in maximum temperature of the warmest month (MTWM) by 2070. Data for RCP6.0.**

**Supplemental Figure S11.**
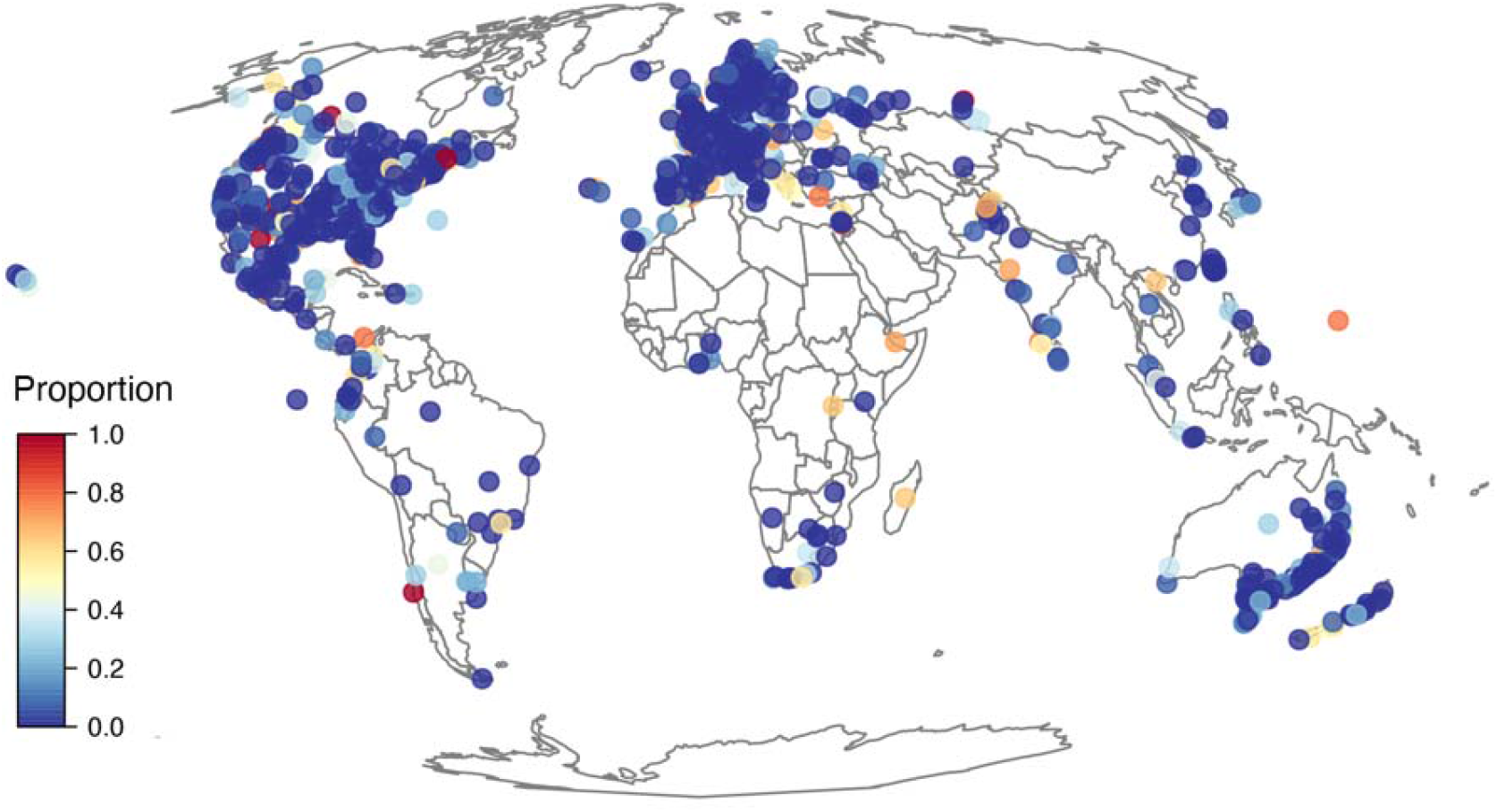
Proportion of urban forest species in each city predicted **to be at risk of future change in precipitation of the driest quarter (PDQ) by 2070. Data for RCP6.0.**

**Supplemental Figure S12.**
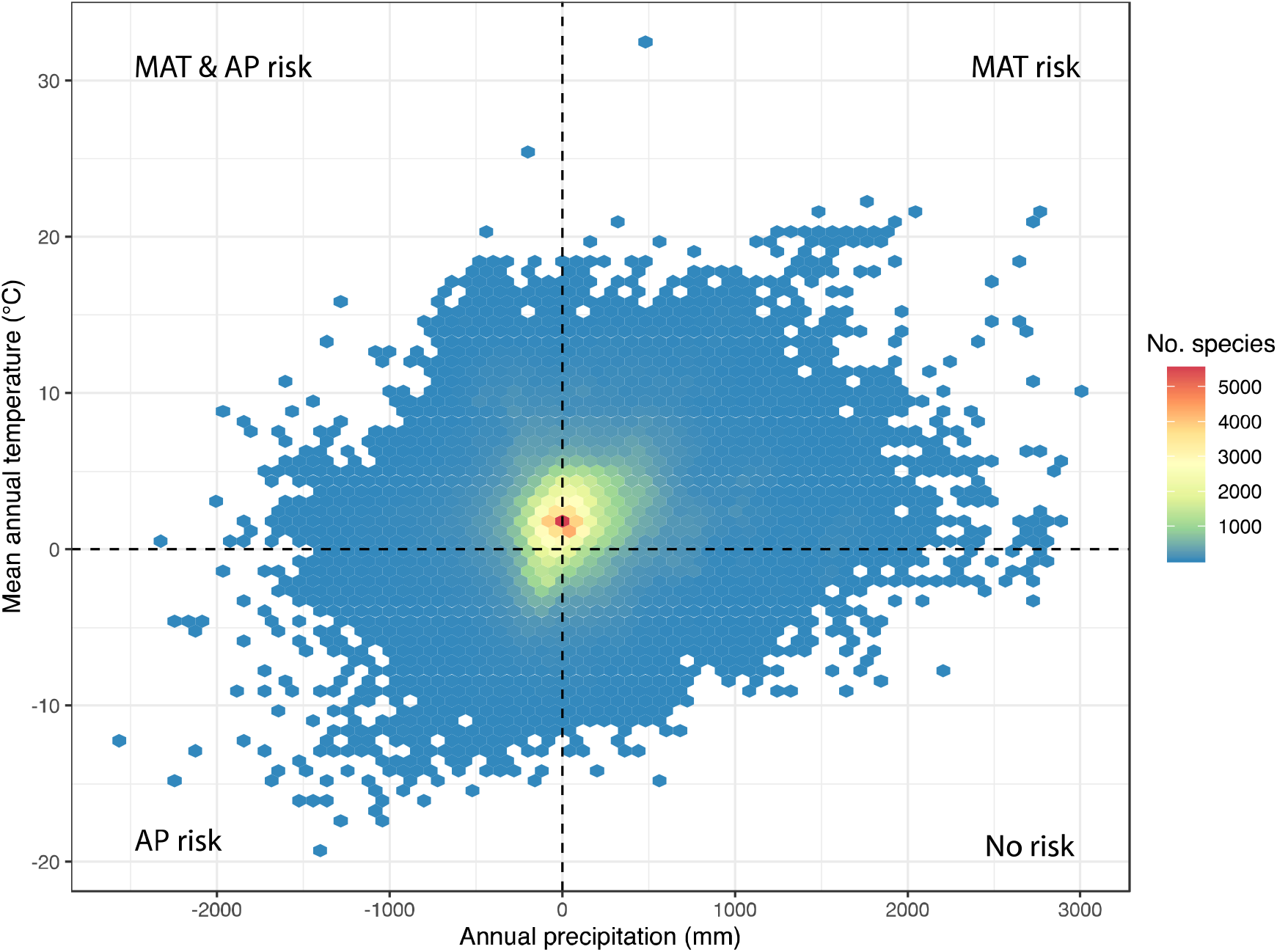
Species at risk of change of changes in mean annual temperature and/or annual precipitation in 1,010 cities in 93 countries. For MAT, *R* > 0 indicates the exposure to future climate is greater than the MAT 95^th^ percentile of the focal species (i.e. high risk), *R* < 0 indicates the exposure to future climate change is still within the range of values allowed by the safety margin (*S*), thus it is “safe” under future conditions (i.e. low risk). For AP, *R* > 0 indicates the exposure to future climate is lower than its current safety margin (i.e. low risk), whereas *R* < 0 indicates high risk, as the exposure (*E*) to future climate change is outside the species’ safety margin. Note that the count of species exceeds the number of species (i.e. 16,006 species), as one species can have a different safety margin depending on the city where it is planted.

**Supplemental Figure S13.**
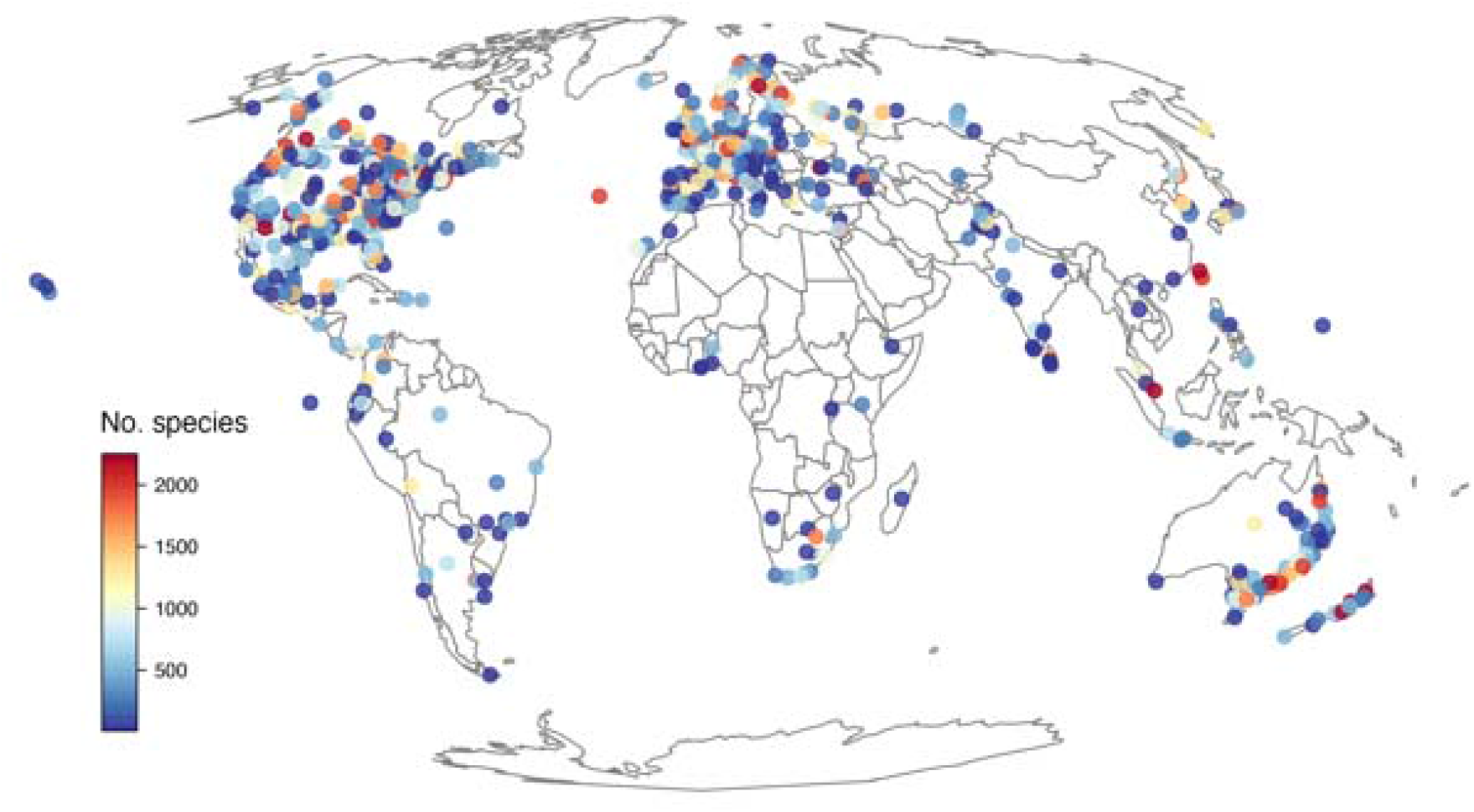
Number of plant species recorded in 1,010 cities.

**Supplemental Figure S14.**
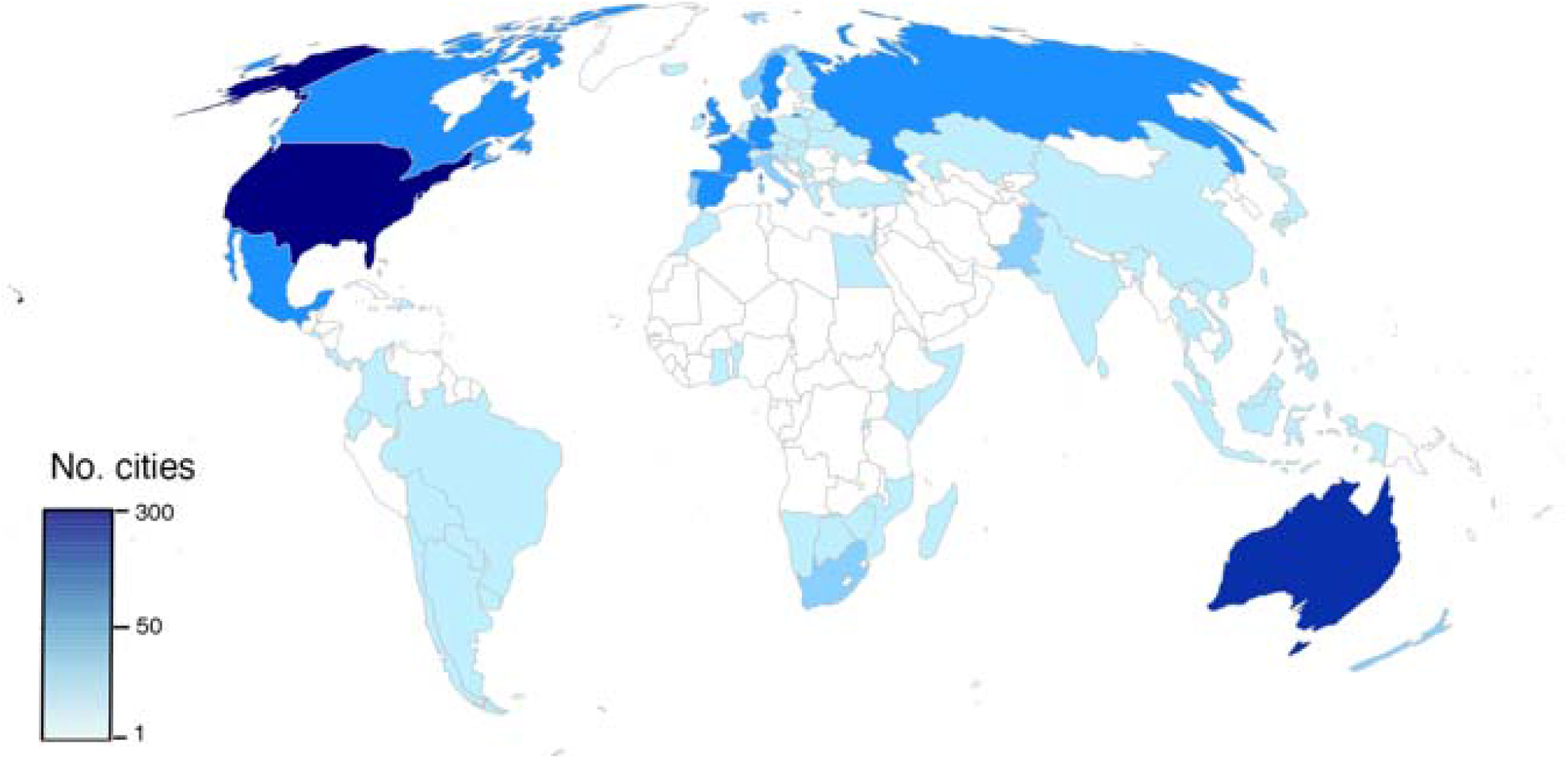
Number of cities assessed in 93 countries.

**Supplemental Figure S15.**
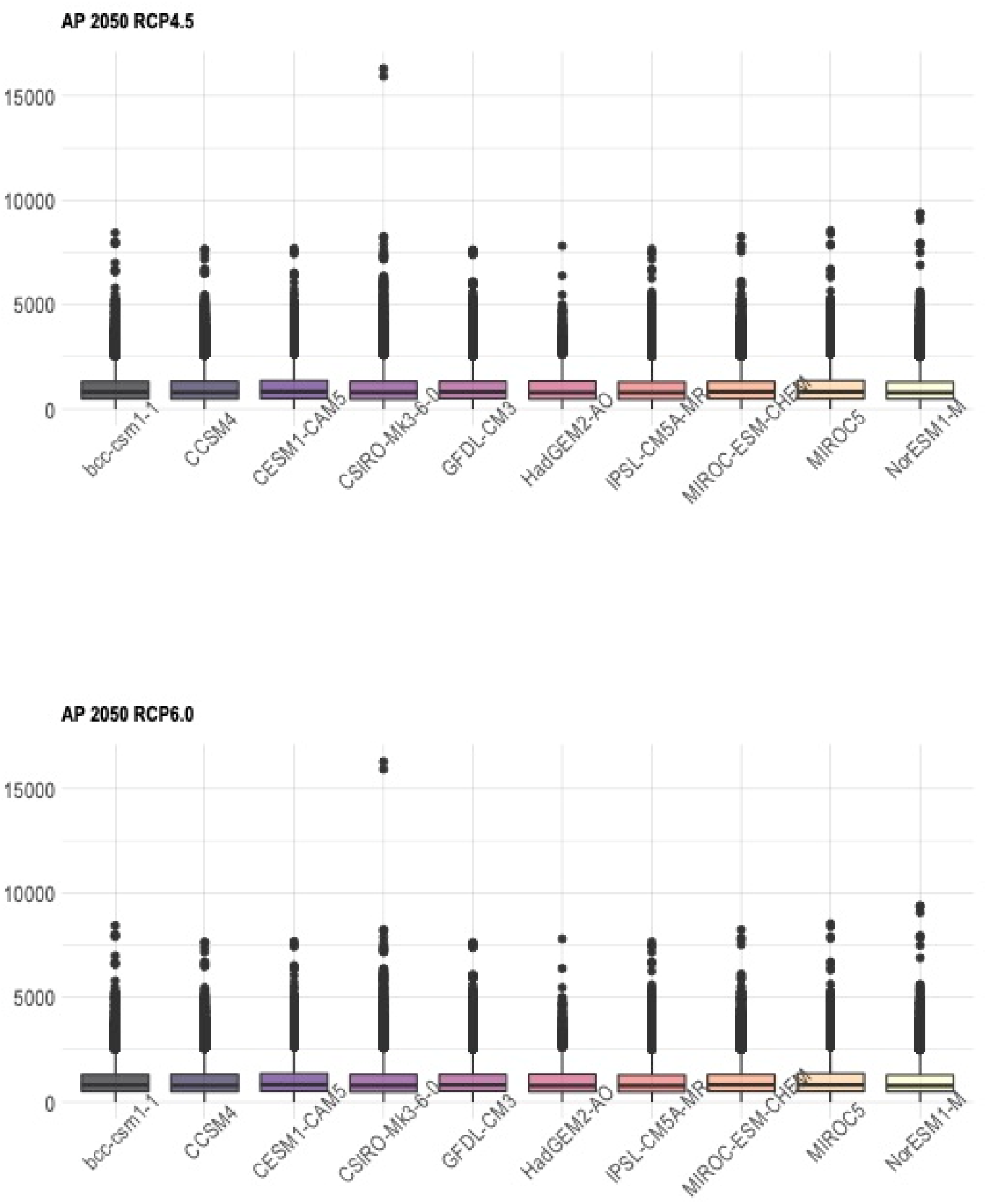
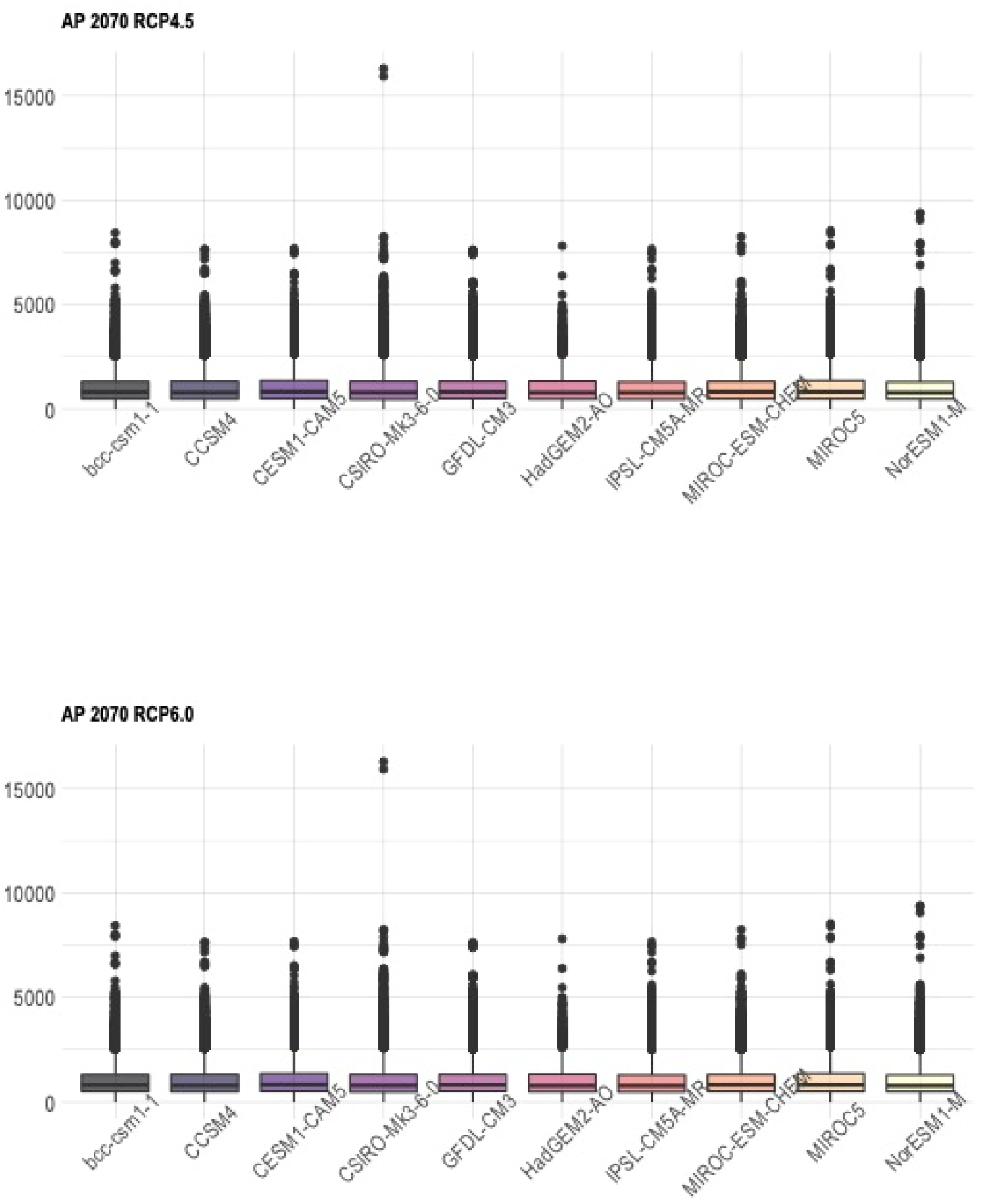
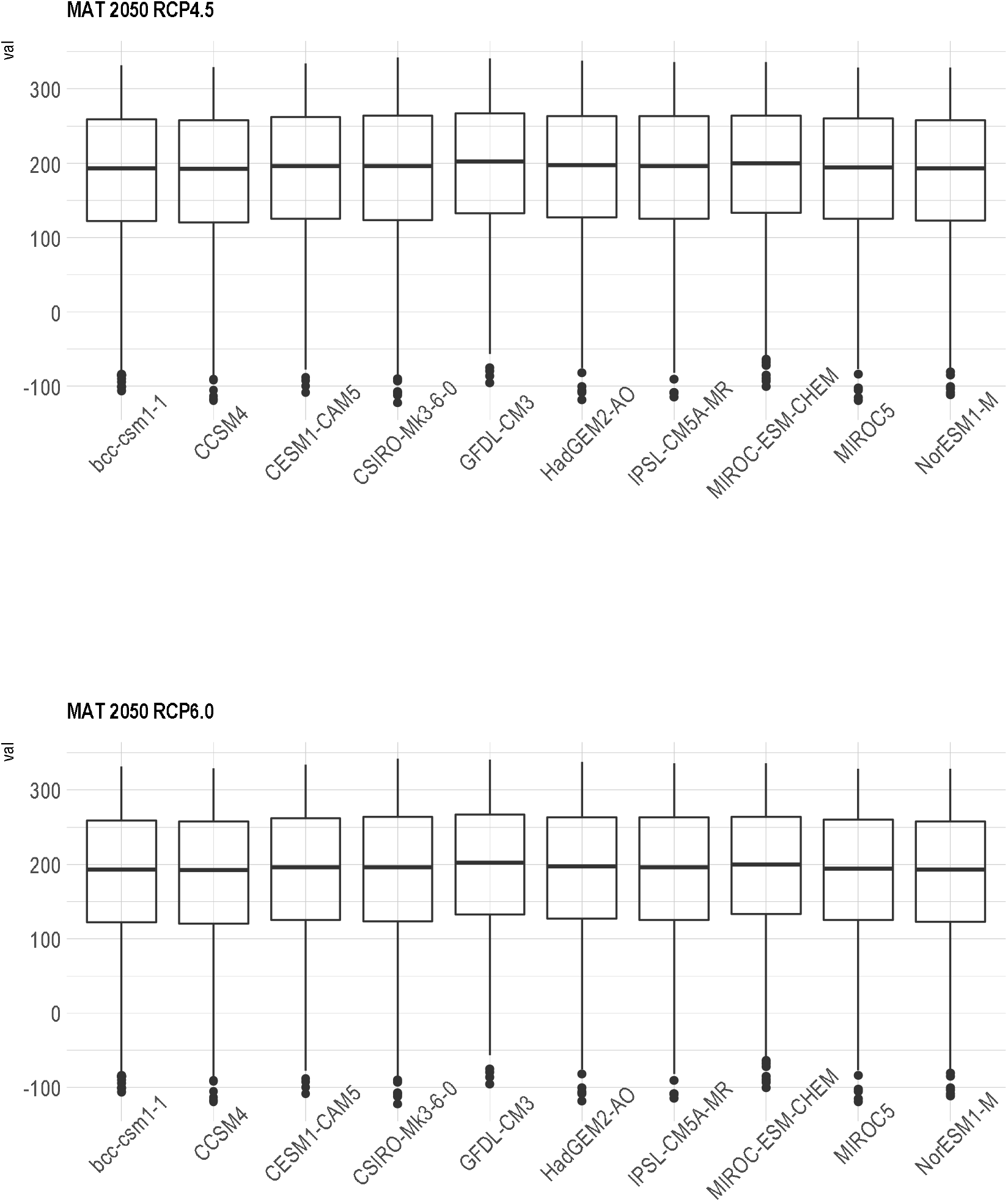
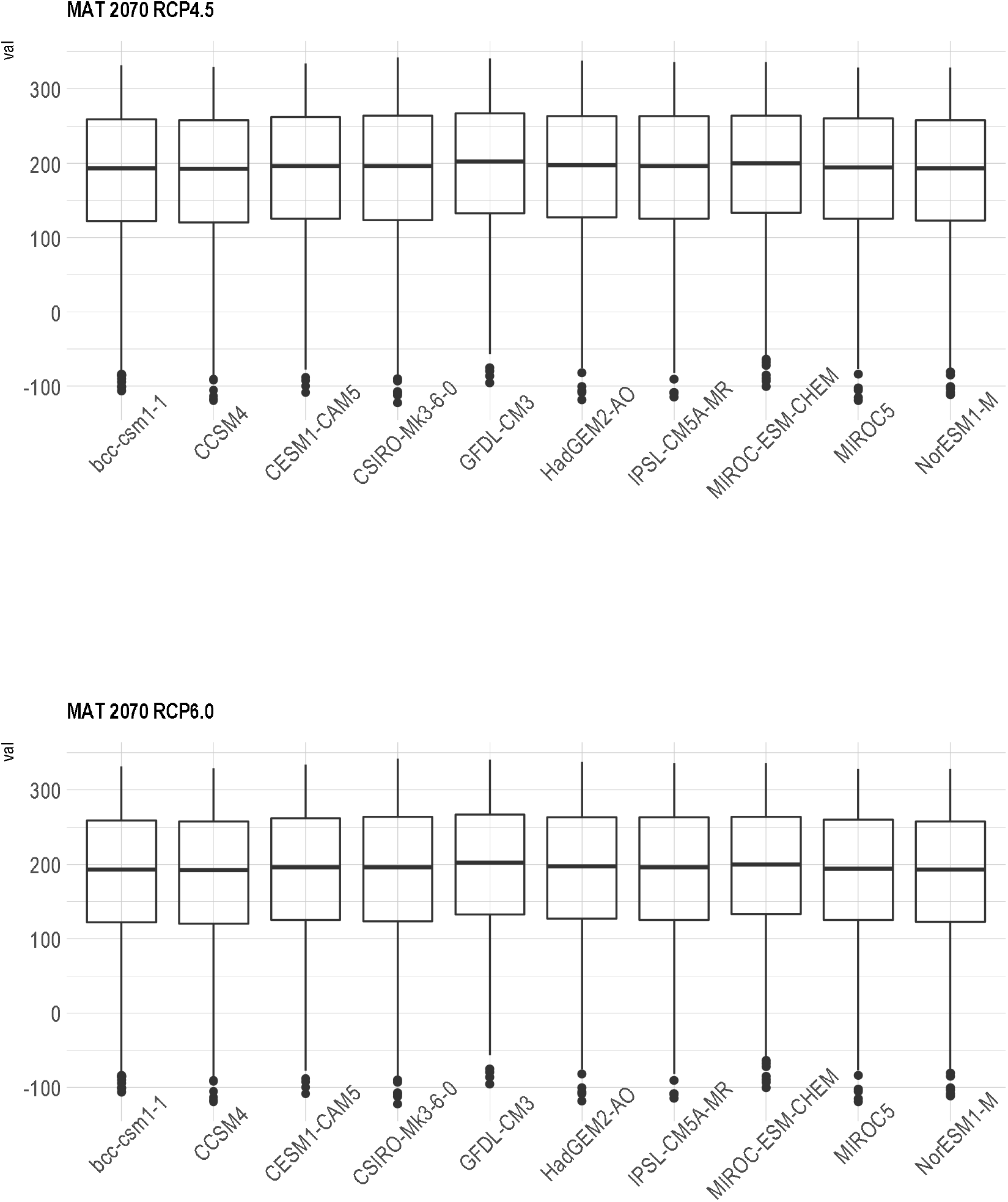
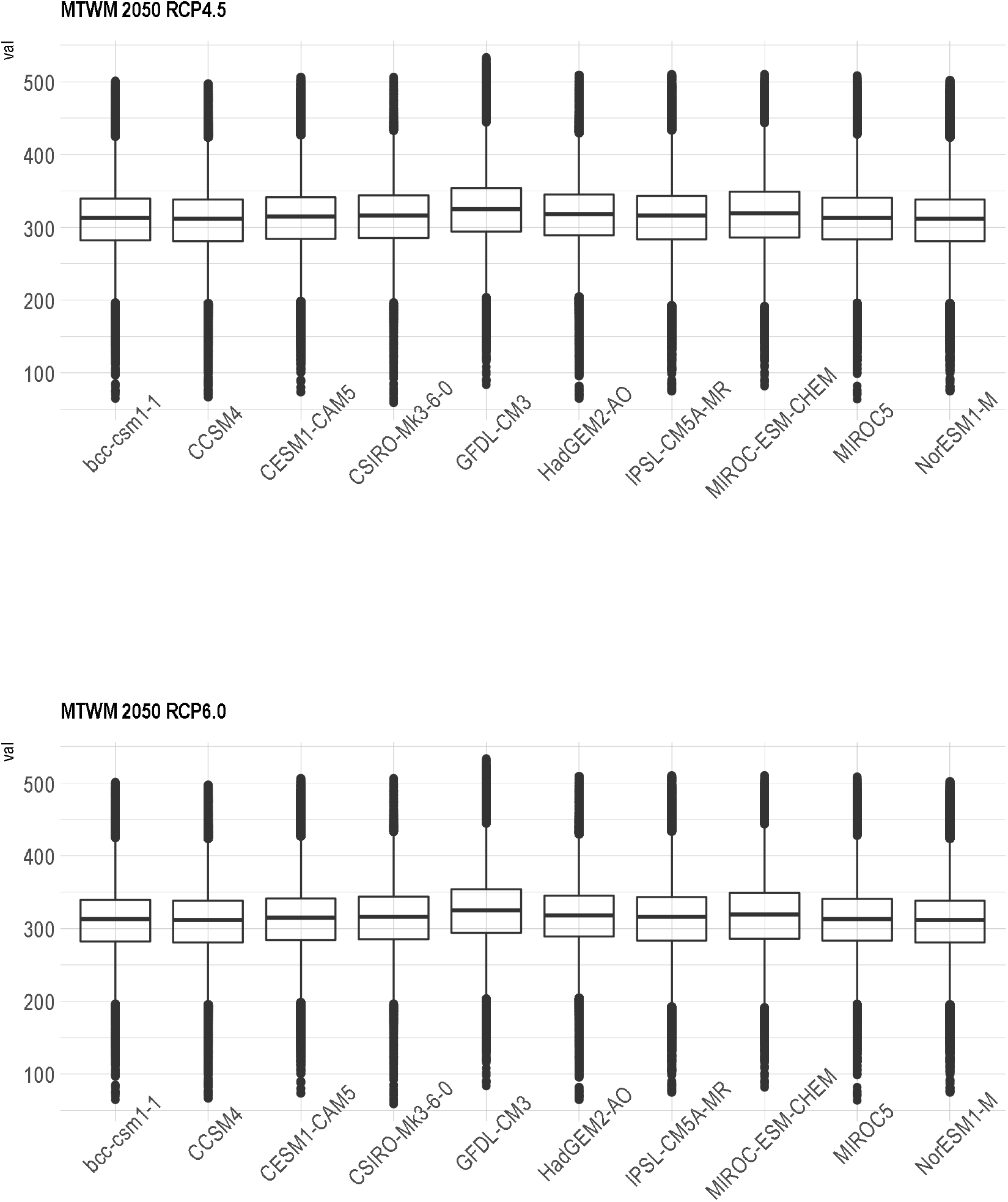
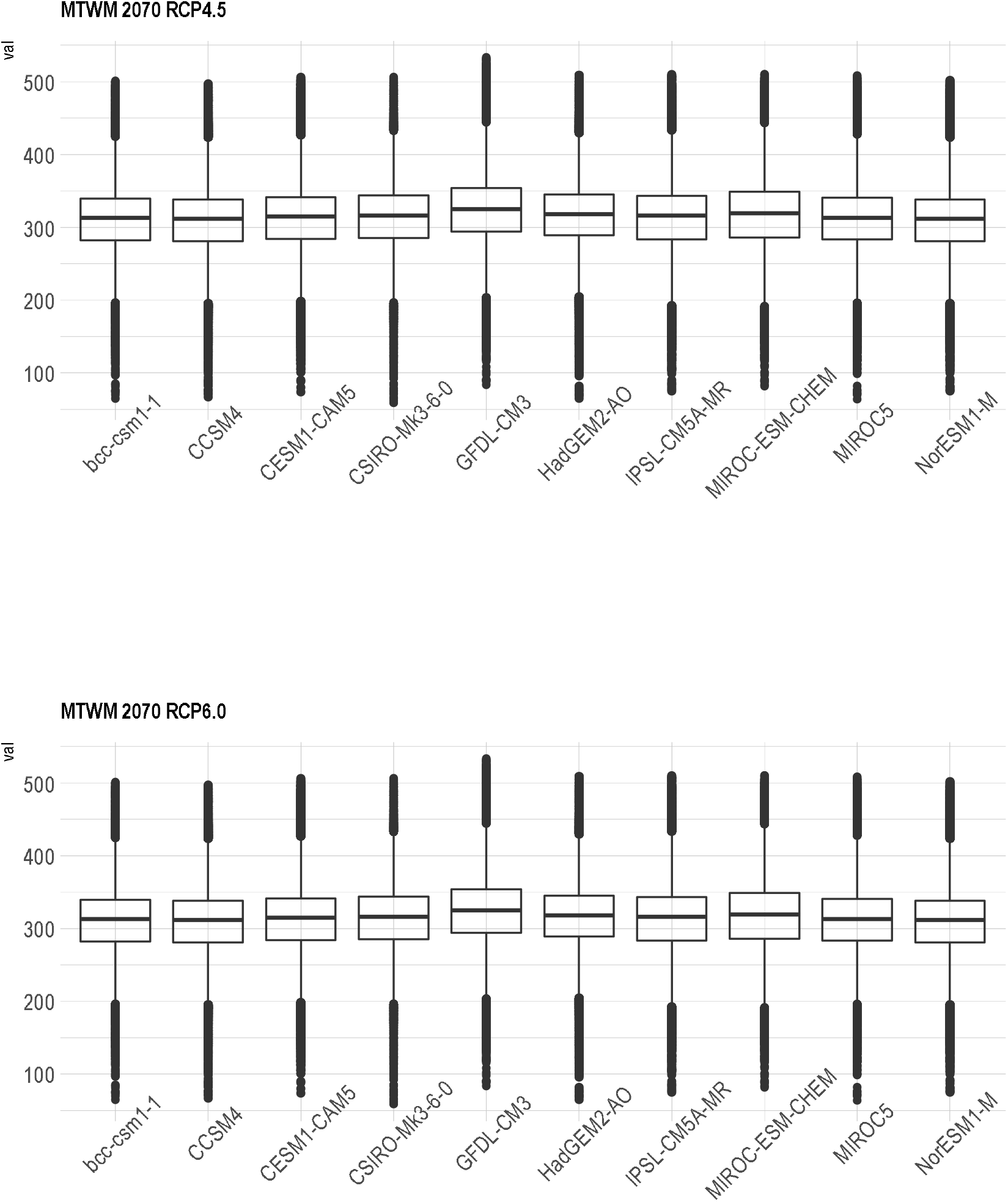
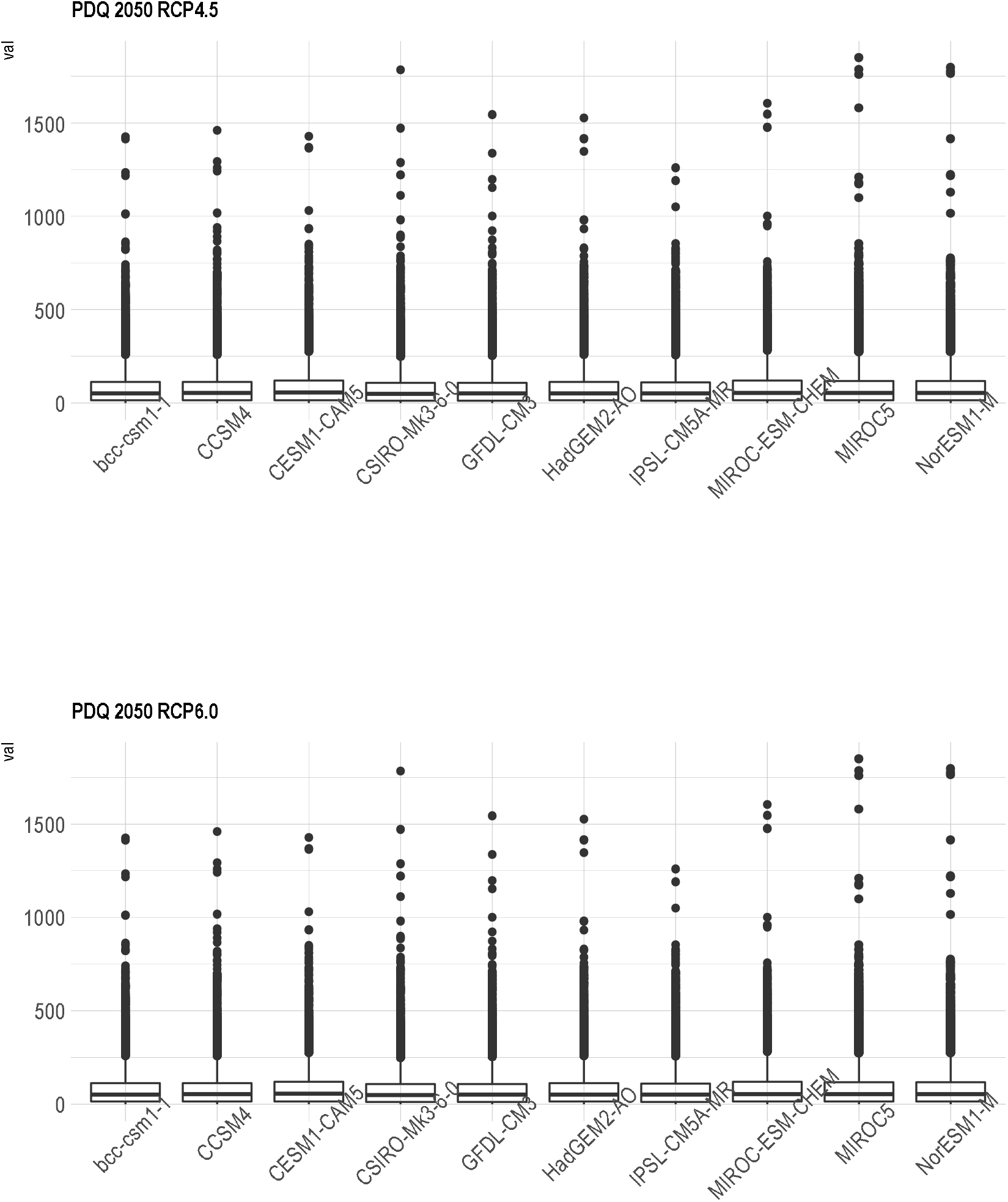
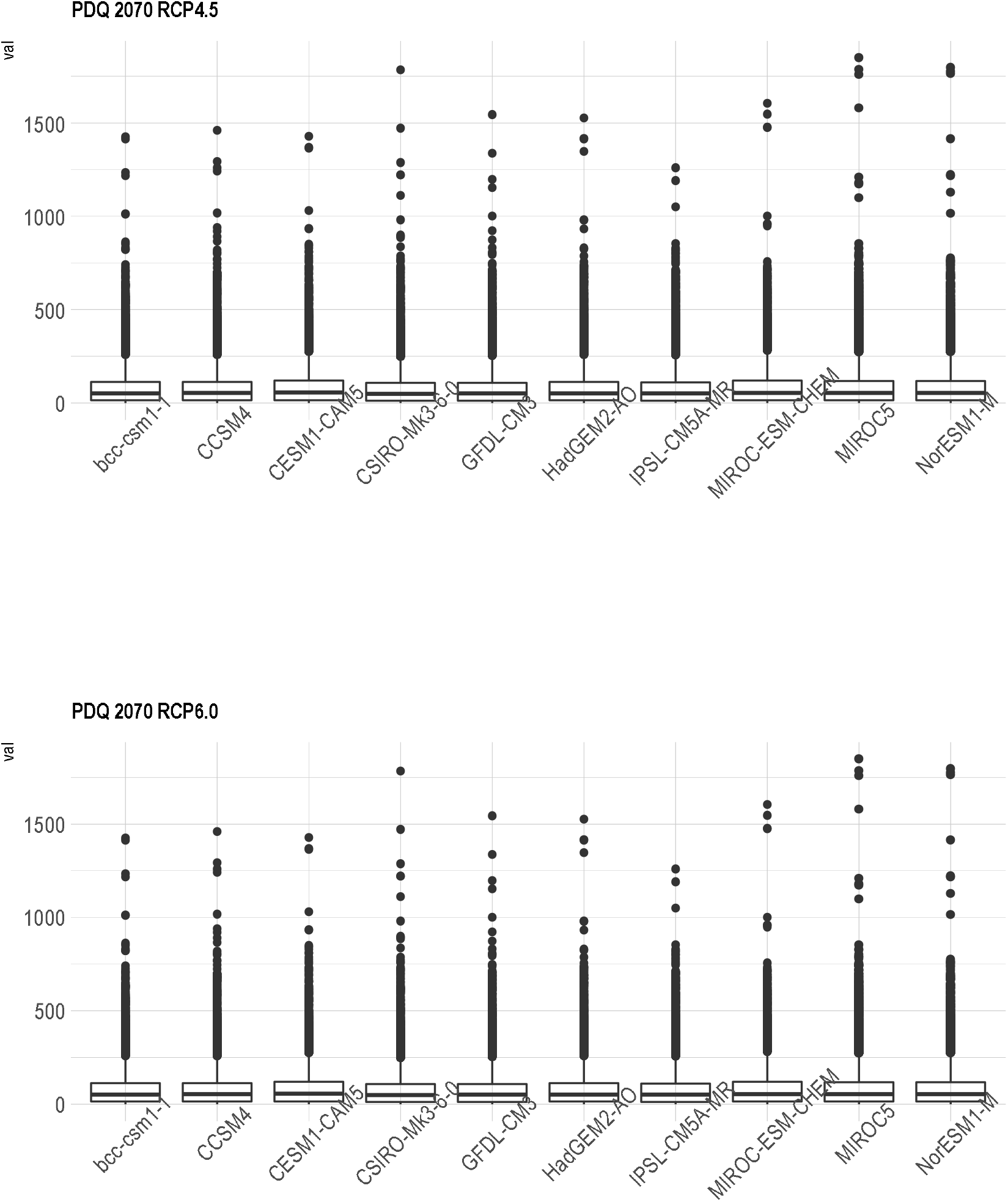
Boxplot showing variability of 10 General Circulation Models (GCMs) for four climatic variables (AP: annual precipitation; MAT: mean annual temperature; MTWM: maximum temperature of the warmer month; and PDQ: precipitation of the driest quarter); two time periods (2050: average for 2041-2060; 2070: average for 2061-2080), and two Representative Concentration Pathway (RCP 4.5 and 6.0).

## Notes

### Competing Interest Statement

The authors have declared no competing interest.

